# Evolution of Phototransduction Genes in Lepidoptera

**DOI:** 10.1101/577031

**Authors:** Aide Macias-Muñoz, Aline G. Rangel Olguin, Adriana D. Briscoe

**Author notes:** Corresponding author: Adriana D. Briscoe. Co-first authors, these authors contributed equally to the work (Aide Macias-Muñoz, Aline G. Rangel Olguin).

## Abstract

Vision is underpinned by phototransduction, a signaling cascade that converts light energy into an electrical signal. Among insects, phototransduction is best understood in *Drosophila melanogaster.* A survey of phototransduction genes in four insect genomes found gains and losses between *D. melanogaster* and other insects; this study did not include lepidopterans. Diurnal butterflies and nocturnal moths occupy different light environments and have distinct eye morphologies, which might impact the expression of their phototransduction genes. Here, we used transcriptomics and phylogenetics to identify phototransduction genes that vary between *D. melanogaster* and Lepidoptera, and between moths and butterflies. Most phototransduction genes were conserved between *D. melanogaster* and Lepidoptera, with some exceptions. We found two lepidopteran opsins lacking a *D. melanogaster* ortholog, and using antibodies found that one, a candidate retinochrome which we name unclassified opsin (UnRh), is expressed in the crystaline cone cells and the pigment cells of the butterfly *Heliconius melpomene*. We also found differences between Lepidoptera and *D. melanogaster* phototransduction in diacylglycerol regulation where a lepidopteran paralog, DAGβ, may be taking on a role in vision. Lastly, butterflies express similar amounts of *trp* and *trpl* channel mRNAs, while moths express approximately 50x less *trp*. Since TRP/TRPL channels allow Ca^2+^ and Na^+^ influx this might explain why moths appear to express less *Calx* and *Nckx30C* Na^+^/Ca^2+^ channel mRNAs. Our findings suggest that while many single-copy *D. melanogaster* phototransduction genes are conserved in lepidopterans, phototransduction gene expression differences exist between moths and butterflies that may be linked to their visual light environment.

## Introduction

Phototransduction is the process that underlies vision, in which light is converted into an electrical signal. Vision has intrigued scientists for many years making phototransduction one of the best-studied signaling pathways (Shichida & Matsuyama 2009). The genes and proteins involved in *D. melanogaster* phototransduction have been investigated for over 40 years (Hardie 2001; Hardie & Raghu 2001; Katz & Minke 2009; Montell 2012; Hardie & Juusola 2015). However, studies of phototransduction cascade genes in other insects are largely lacking. A comparison of vision-related genes in four insect genomes (mosquito, red flour beetle, honeybee and fruit fly) found gains and losses of genes involved in phototransduction across different lineages (Bao & Friedrich 2009). *D. melanogaster* had by far the largest number of gene gains compared to the other insects examined. Other insect species might also differ in the genes underlying phototransduction.

Phototransduction takes place in specialized neurons known as photoreceptor cells whose microvilli incorporate light-sensitive opsin proteins bound to a retinal-derived molecule called a chromophore (Fain et al. 2010). Phototransduction begins when light is absorbed by the chromophore (11-*cis*-3-hydroxyretinal in *D. melanogaster*) causing the chromophore to change its confirmation from *cis* to all-*trans* (von Lintig et al. 2010). In *D. melanogaster*, this change in configuration triggers a G-protein-coupled cascade (similar to Figure 1) that activates phospholipase C (PLC) (Bloomquist et al. 1988). PLC hydrolyses phosphatidylinositol 4,5-bisphosphate (PIP_2_) to produce inositol 1,4,5-trisphosphate (InsP_3_) and diacylglycerol (DAG) (Bloomquist et al. 1988; Hardie 2001). Concurrently, by a mechanism that is not well understood, there is an opening of Ca^2+^-permeable light-sensitive transient receptor potential (TRP) and transient receptor potential-like (TRPL) channels which causes depolarization of the cell (Montell & Rubin 1989; Hardie & Minke 1992; Niemeyer et al. 1996; Shieh & Zhu 1996; Montell 2005). Finally, phototransduction is terminated when the activated rhodopsin (metarhodopsin) binds arrestin (Dolph et al. 1993; Stavenga & Hardie 2011).

**Figure 1:**
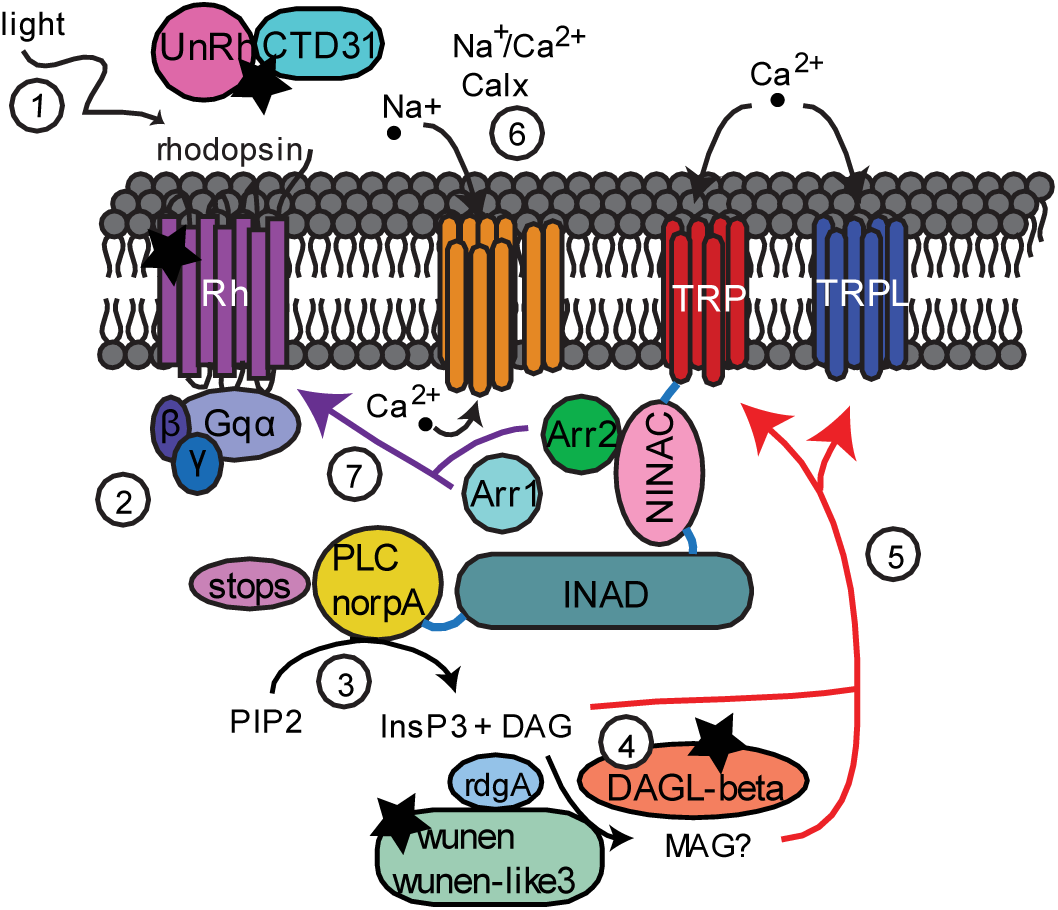
Speculative model of the lepidopteran phototransduction cascade. 1) Light activates rhodopsin by a configurational change of the chromophore from 11-*cis* to all-*trans*. The chromophore is transported by Hme CTD31 and photoisomerized from all-*trans* to 11-*cis* by the unclassified opsin. 2) G_q_*α* is released from a G-protein complex of 3 subunits (*α, β*, and *γ*) and activates phospholipase C (PLC). 3) PLC hydrolyses phosphatidylinositol 4,5-bisphosphate (PIP_2_) to produce inositol 1,4,5-trisphosphate (InsP_3_) and diacylglycerol (DAG). 4) Diacylglycerol lipase (DAGL-beta) hydrolyzes DAG to produce monoacylglycerol (MAG). 5) DAG and MAG may activate TRP and TRPL, by a mechanism that has not been established. 6) Na^+^/Ca^2+^ exchanger channel pumps Ca^2+^ out of the photoreceptor cell. 7) Arrestin 1 and 2 bind rhodopsin to terminate the cascade with Arrestin 2 being the dominant arrestin in both *D. melanogaster* and butterflies. Stars signify differences in phototransduction between *D. melanogaster* and Lepidoptera, either duplication or upregulation of vision-related gene paralogs in butterfly heads.

Numerous studies have focused on characterizing the opsins expressed in photoreceptor cells and their arrangement across the compound eye (Spaethe & Briscoe 2005; Henze et al. 2012; Futahashi et al. 2015; McCulloch et al. 2016; Perry et al. 2016; Giraldo-Calderón et al. 2017; McCulloch et al. 2017). Although a large focus is on the opsins, changes in the downstream pathway by which opsins function might also contribute to differences in visual systems (Plachetzki et al. 2010). Fewer studies have investigated the downstream phototransduction cascade in non-*D. melanogaster* insects. Studies of phototransduction in other insects have focused on presence, absence, or relative expression of genes in head transcriptomes. In the troglobiont beetle, *Ptomaphagus hirtus*, for example, 20 genes were identified from adult head mRNA (Friedrich et al. 2011). Exposure of the oriental armyworm, *Mythimna separate*, to different light environments resulted in differential expression of phototransduction genes in adult heads (Duan et al. 2017). Similarly, phototransduction genes were also differentially expressed between seasonal forms in heads of the butterfly *Bicyclus anynana* (Macias-Muñoz et al. 2016). One study quantified opsin and TRP channel gene expression and used RNAi to determine that opsin has the largest effect on phototransduction in the nocturnal cockroach *Periplaneta americana* (but see below)(French et al. 2015). Yet, it remains largely unknown how variable the phototransduction cascade is between insect species.

Lepidoptera, moths and butterflies, provide an interesting group in which to investigate the molecular evolution and expression of phototransduction genes in insects adapted to different light environments (Yagi & Koyama 1963; Horridge et al. 1972; Nilsson et al. 1984; Yack et al. 2007; Warrant & Dacke 2016). Phylogenetic analyses have been used to reveal duplications of opsin genes that had not been previously described (Spaethe & Briscoe 2004; Sison-Mangus et al. 2008; Briscoe et al. 2010). A survey of 23 vision-related gene families in 19 metazoan genomes revealed that eye development and phototransduction genes have higher rates of retention and duplications in pancrustaceans (Rivera et al. 2010). Since only the nocturnal domesticated silkmoth *Bombyx mori* was used in the pancrustacean study and only five gene families involved in phototransduction were examined (r-opsin, TRP, phospholipase C, Gq-alpha and arrestin) (Rivera et al. 2010), it remains to be seen if there are additional differences in phototransduction genes between *D. melanogaster* and moth and butterfly species. In our present study, we expand on the genes surveyed thus far by looking at 76 phototransduction-related genes. Phylogenetic analyses of phototransduction genes in Lepidoptera may reveal: 1) the extent to which *D. melanogaster* phototransduction genes are duplicated or deleted in Lepidoptera, 2) lepidopteran-specific phototransduction features, and 3) differences between diurnal and nocturnal Lepidoptera.

While gene trees tell the probable evolutionary history of gene families, gene expression data is a step towards inferring gene function. Genes involved in vision should be highly expressed in photoreceptor cells and upregulated in the eyes relative to other tissue types, thus visualizing or quantifying where phototransduction genes are expressed will reveal whether they have a potential role in vision. As an example, the horseshoe crab *Limulus polyphemus* has 18 opsins, some of which are expressed only in the eyes, in eyes and central nervous system, exclusively in the central nervous, and some are not expressed in either (Battelle et al. 2016). It is possible that the opsins missing from the eyes and central nervous system are expressed in other tissue types and have non-visual functions (Feuda et al. 2016) or are not expressed at all. Similarly, the butterfly *Heliconius melpomene*, reference genome (Davey et al. 2016) revealed a *UVRh* duplication but mRNA shows that one of the copies is downregulated in this species and only one of the copies has protein expression in the compound eye (McCulloch et al. 2017). Studies such as this highlight the importance of quantifying gene expression in candidate tissues before inferring gene function based on sequence alone. Further, it is also possible that a paralogous member of the same gene family in fact partakes in the predicted visual function. As an example, an expression analysis of CRAL-TRIO domain genes in *H. melpomene* found that an ortholog of *D. melanogaster pinta* is missing in Lepidoptera (Wang & Montell 2005, Smith and Briscoe 2015). Instead a lepidopteran paralog appears to carry out a similar function of chromophore binding (Macias-Muñoz et al. 2017). Moreover, as observed in the cockroach, while genes such as *TRP* and *TRPL* are conserved and expressed, one gene copy (*TRPL*) might have a greater impact on phototransduction than the other (French et al. 2015). Consequently, investigating both gene gain/loss and the expression of phototransduction genes in Lepidoptera might uncover differences in their visual processing that helps them function in different light environments.

In this study, we combined transcriptomics and phylogenetics to perform the first investigation of candidate phototransduction genes in Lepidoptera. We used RNA-Sequencing data from four tissues of the butterfly *Heliconius melpomene* to identify genes upregulated in heads. We hypothesized that genes upregulated in heads, might have eye and vision-related functions. A functional enrichment analysis suggested that many of the genes upregulated in *H. melpomene* heads function in phototransduction. To investigate the molecular evolution and to identify gene gain or loss between *D. melanogaster* and Lepidoptera and between moths and butterflies, we extracted 76 phototransduction gene sequences from reference genomes of eight insect species including the moth, *Manduca sexta*, and the butterflies, *Danaus plexippus* and *Heliconius melpomene* (Zhan et al. 2011; Davey et al. 2016; Kanost et al. 2016). Then we generated 32 phylogenetic trees. In case any genes were missing annotations in the reference assemblies, we searched *de novo* transcriptome assemblies from *M. sexta, H. melpomene*, and *D. plexippus*. We found that most of the phototransduction pathway is conserved between Lepidoptera and *D. melanogaster*, with some exceptions (Figure 1). Our methods allowed us to uncover two lepidopteran opsin genes that lack a homolog in *D. melanogaster*, one of which we verified using antibodies, is expressed in pigment cells. In addition, diacylglycerol regulation appears to differ between Lepidoptera and *D. melanogaster*, where a paralogous gene in lepidopterans, *DAGβ*, maybe taking on a role of a lost ortholog of *D. melanogaster DAGα*. While we found no copy-number differences between moths and butterflies in the investigated phototransduction genes, we discovered an important difference between moths and butterflies in their expression of vision-related ion channels, *trp, Calx*, and *Nckx30C*.

## Materials and Methods

### Transcriptome-wide differential expression analysis

RNA-Sequencing data for *H. melpomene* male and female head, antennae, legs and mouth parts were obtained from array express projects E-MTAB-1500 and E-MTAB-6249 (Table S1). A four tissue *de novo* transcriptome made from one library per tissue type per sex was used as reference (doi:10.5061/dryad.857n9; see Macias-Muñoz et al. 2017). Reads from each sample were mapped to the transcriptome using bwa (Li & Durbin 2009) and RSEM (Li & Dewey 2011) was used to quantify mapped raw reads. We used edgeR (Robinson et al. 2010) to perform three pairwise comparisons for differential expression analysis: head versus antennae, head versus legs, and head versus mouth parts. For each comparison, a generalized linear model was used to include terms for batch, tissue, sex, the interaction of sex and tissue (∼batch + tissue + sex + sex*tissue). Each analysis also included filtering to remove lowly expressed contigs (less than 1 count per million for at least 4 groups). Samples were normalized using a trimmed mean of the log expression ratios (TMM) (Robinson & Oshlack 2010). After each comparison, p-values were further corrected using a Bonferroni false discovery rate (FDR) correction. Contigs were considered significantly differentially expressed when the FDR was less than 0.05 and the log fold change (logFC) was greater than 1.

Of these differentially expressed contigs, we identified which were upregulated in heads for each comparison. The resulting gene lists were merged to identify contigs commonly upregulated in heads. Patterns of expression for significant contigs and those commonly upregulated in heads were visualized using heatmaps (Ploner 2012). Contigs were annotated with *D. melanogaster* gene IDs (Marygold et al. 2013) by using command-line BLAST+ to compare *H. melpomene* transcriptome sequences to *D. melanogaster* gene sequences (Camacho et al. 2009). We used batch download in Flybase to acquire gene ontology terms (GO terms) for our differentially expressed and head upregulated contigs. Differentially expressed contigs with unique annotations were enriched for function using a Database for Annotation, Visualization, and Integrated Discovery (DAVID) (Huang et al. 2009). Contigs commonly upregulated in heads were also assigned GO terms and protein classification NCBI Blast and InterProScan in BLAST2GO to uncover additional annotations potentially missing from a comparison to *D. melanogaster* only (Conesa & Götz 2008; Conesa et al. 2005; Götz et al. 2008).

### Phototransduction genes in insect genomes

To identify phototransduction genes in Lepidoptera and explore their evolutionary history, we used *D. melanogaster* sequences to search for homologs in published insect genomes. We began with a compilation of sequences by Bao and Friedrich (Bao & Friedrich 2009) but expanded it to include Lepidoptera species and additional phototransduction genes (Table S2). We used BLAST to search the genomes of *Anopheles gambiae, Apis mellifera, Tribolium casteum, Bombyx mori, Manduca sexta, Heliconius melpomene*, and *Danaus plexippus*. Sequences with identity of more than 20% and an e-value greater than 1E-10 were tested for homology using reciprocal blastp to the NCBI database. In addition to searching Lepidoptera reference genomes, we searched *de novo* transcriptome assemblies to improve annotation and find duplicates that are not found in genomes. We searched an *H. melpomene* four tissue transcriptome (doi:10.5061/dryad.857n9; Macias-Muñoz et al. 2017) and a *M. sexta* head transcriptome (doi:10.5061/dryad.gb135; Smith et al. 2014). The nucleotide sequences recovered from *de novo* transcriptomes were translated using OrfPredictor with the blastx option before testing them by reciprocal blast hits (Min et al. 2005).

Sequence corrections were accomplished by aligning sequences in Molecular Evolutionary Genetics Analysis (MEGA) and manually correcting missing pieces then using BLAST to recover the segment from the genome. To obtain the consensus sequences, we inputted corrected sequences to CLC Genomics (CLCBio) and mapped reads against them. With some exceptions, we recovered the entire sequence for all phototransduction genes in *H. melpomene* (Table S3) and *M. sexta* (Table S4). Phototransduction genes for *H. melpomene* and *M. sexta* were annotated and deposited in GenBank with accession numbers XXXXXXXX-XXXXXXXX (Table S5). We also searched NCBI for the insect sequence matches to our annotated transcriptome sequences, and added any hits missing from our data set. In addition, to test the history of *inaE* gene in *D. melanogaster* and the lepidopteran *DAGLβ-like* in the context of the evolution of the gene family in animals, we added *Homo sapiens, Mus musculus*, and *Hydra vulgaris* homolog sequences using blastp searches of NCBI data bases for *H. sapiens* (taxid:9606), *M. musculus* (taxid:10090), *H. vulgaris* (taxid:12836).

Protein sequences for each gene family were aligned in MEGA 7.0 using the Multiple Sequence Comparison by Log-Expectation (MUSCLE) (Edgar 2004; Kumar et al. 2016). The alignments were further corrected manually. Before generating maximum likelihood trees, we calculated Bayesian Information Criterion (BIC) values to assess which substitution model would best fit our data (Schwarz 1978; Kumar et al. 2016). We used the best fit model to generate phylogenies using 100 bootstrap replicates (Table S6).

### Expression of candidate genes

To study expression patterns among homologs, we looked at the expression of genes belonging to five gene families in *M. sexta* heads and in *H. melpomene* heads, antennae, legs, and mouth parts (labial palps + proboscis). Rearing conditions for *M. sexta* are described in Smith et al. (2014) and for *H. melpomene* in Briscoe et al. (2013) and Macias-Muñoz et al. (2017). We began by adding our corrected *H. melpomene* and *M. sexta* sequences (Table S3-5) to the *de novo* transcriptome assembly. We uniquely mapped trimmed and parsed reads from four male and four female *M. sexta* heads (E-MTAB-2066; Smith et al. 2014) to the corrected *M. sexta* transcriptome using bowtie (Langmead et al. 2009). We also mapped processed reads from *H. melpomene* head, antennae, legs and mouth parts (E-MTAB-1500, E-MTAB-6249, E-MTAB-6342; Macias-Muñoz et al. 2017) to the corrected *H. melpomene* transcriptome. RSEM was used to count raw reads mapped (Li & Dewey 2011). We visualized expression levels by graphing Transcripts Per Million (TPM) for each gene of interest using ggplot2 (Wickham 2009). Differential expression between tissue types for *H. melpomene* was repeated as outlined above in edgeR using uniquely mapped reads to transcriptome with corrected sequences. However for this data set to allow for less stringency, we used qvalue (Dabney & Storey 2013) to correct p-values rather than Bonferroni.

### Immunohistochemistry

An antibody was generated against the motif N-CKGARTVDEDKKKE-C of the *H*. *melpomene* unclassified opsin (UnRh) in guinea pig and was immunoaffinity purified (New England Peptide, Gardner, MA). We also used an antibody against the long-wavelength sensitive opsin (LWRh) of *Limenitis astyanax* (Frentiu et al. 2007) which labels LWRh expressing cells in *Heliconius* (McCulloch et al. 2016). Eyes were fixed, sucrose protected, cryosectioned and immunolabeled according to methods in McCulloch et al. (2016). Following washes with PBS and block (McCulloch et al. 2016; Macias-Muñoz et al. 2017), slides were incubated with 1:15 rabbit anti-LWRh and 1:30 guinea pig anti-UnRh antibodies in blocking solution overnight at 4°C. After washing in PBS, slides were incubated with 1:500 goat anti-rabbit Alexafluor 555 and 1:250 goat anti-guinea pig Alexafluor 633 secondary antibodies in blocking solution for two hours at room temperature in the dark. Slides were washed once more in PBS in the dark and stored for imaging with Aqua Poly/Mount (Polysciences, Inc. Cat. # 18606). Images were at the UC Irvine Optical Biology Core Facility using a Zeiss LSM700 confocal microscope under 20x objective. Two-channel composites were generated using Fiji and brightness was adjusted for clarity using Adobe Photoshop.

## Results and Discussion

### Transcriptome-wide differential expression analysis

To determine the predicted functions of genes expressed in butterfly heads, we used *H. melpomene* RNA-Seq data to identify contigs upregulated in head tissue relative to antennae, legs, and mouth parts. A multidimensional scaling (MDS) plot showed that head libraries group together and away from other tissue types (Figure 2A). Differential expression analysis comparing heads vs. antennae yielded 1,173 Differentially Expressed (DE) contigs (Figure S1; Table S7), 561 of these were upregulated in heads (Table 1). Analysis of head vs. legs mRNA gave 1,472 DE contigs (Figure S1; Table S8), of these contigs 928 were upregulated in heads. Heads vs. mouth parts comparison yielded 1,486 DE contigs (Figure S1; Table S9), 914 of these were upregulated in heads (Table 1). DE contigs from each of the three pairwise comparisons matched 590, 748, and 700 unique gene ontology terms (GO terms; Table 1).

**Table 1.** Summary of *Heliconius melpomene* transcriptome-wide analysis

|  |  |  | Upregulated | Commonly<br>upregulated | Unique GO |
| --- | --- | --- | --- | --- | --- |
|  | qvalue | Bonferroni | in Heads | in heads | terms |
| Head vs. Antennae | 4,868 | 1,173 | 561 |  | 590<sup>†</sup> |
| Head vs. Legs | 6,108 | 1,472 | 928 |  | 748<sup>†</sup> |
| Head vs. Mouth | 6,176 | 1,486 | 914 |  | 700<sup>†</sup> |
| Merged* |  |  |  | 281 | 154 |

\*Merged are genes commonly upregulated in heads after merging results of pairwise comparisons.

**Figure 2:**
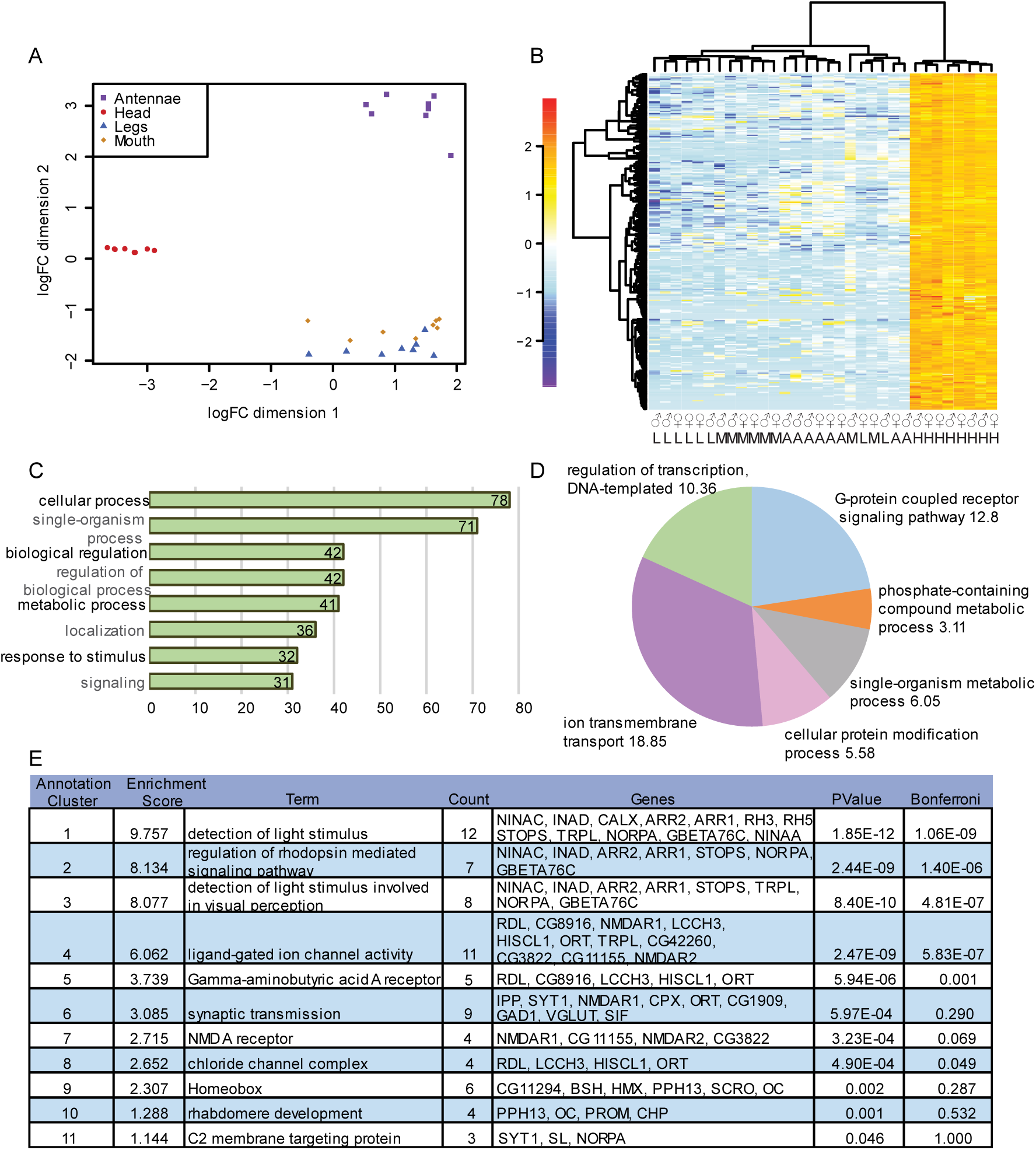
Differential expression analysis. (A) Multidimensional scaling (MDS) plot of RNA-seq libraries from *H. melpomene* antennae, head, legs and mouth parts. (B) Heatmap of genes commonly upregulated in heads, numbers indicate log-fold change. (C) Level 2 biological process terms for genes commonly upregulated in heads using Blast2GO. (D) Multilevel pie chart summary of GO terms with node score information using Blast2GO. A node score is the number of sequences associated to with a particular GO term. (E) Enrichment results for genes commonly upregulated in heads and annotated using FlyBase and DAVID.

We performed functional enrichment analyses for contigs DE between the 3 comparisons (head vs. legs, head vs. antennae, and head vs. mouth parts) to investigate the potential functions of these genes. We found that DE contigs for the three comparisons had some similar annotation clusters. Annotation terms that were similar across the three comparisons included detection of light stimulus, regulation of rhodopsin mediated signaling, and homeobox domain (Table S10). The first two annotation clusters included genes involved in phototransduction. The homeobox cluster included genes involved in antennal, leg and neuron development, as well as genes involved in compound eye development and morphogenesis such as *araucan, PvuII-PstI homology 13, ocelliless*, and *eyegone*. An annotation term unique to the head vs. antennae comparison was glucose-methanol-choline oxidoreductase which included the genes *glucose dehydrogenase* and *ninaG* among other yet unnamed genes (Table S10). An annotation term unique to the head vs. mouth parts comparison was ion channel activity and included genes involved in perception of touch, taste, and olfaction (Table S8). This cluster also included genes potentially involved in phototransduction such as *cacophony, NMDA receptor 1*, and *transient receptor potential-like* (*trpl*; Table S10).

Most of the genes enriched in the DE analyses between heads and other tissues are biased towards vision, as has also been found in a recent transcriptomic analysis of *M. sexta* adult head tissue alone (Smith et al. 2014). This could be because more transcription is actively occurring in the adult butterfly head and the head is mostly composed by the eye and optic lobe (Girardot et al. 2006). *Heliconius* butterflies have large eyes due to selective pressures that favor development of large eyes regardless of body size and the optic lobe accounts for approximately 64% of the total brain volume (Seymoure et al. 2015; Montgomery et al. 2016).

### Head upregulated genes

We merged the lists of contigs upregulated in heads in each pairwise comparison to obtain 281 contigs commonly upregulated in heads across the three comparisons (Figure 2B; Table 1). Head upregulated contigs annotated using BLAST2GO level 2 analysis showed that 78 of the annotated genes were involved in cellular processes and 32 were involved in response to stimulus (Figure 2C) (Götz et al. 2008; Conesa & Götz 2008; Conesa et al. 2005; Götz et al. 2011). A breakdown of these genes shows that a majority are involved in ion transmembrane transport and G protein coupled receptor signaling pathway (Figure 2D).

Commonly upregulated contigs in heads across the three comparisons corresponded to 154 unique *D. melanogaster* GO terms (Table 1; Table S11). These contigs were grouped into nine annotation clusters using the highest stringency in DAVID (Figure 2E; Huang et al. 2009). The top three annotation clusters were: 1) detection of light stimulus, 2) regulation of rhodopsin-mediated signaling pathway and 3) detection of light stimulus involved in visual perception (Figure 2E). The genes grouped within these clusters were annotated with phototransduction functions due to possible homology to *D. melanogaster* genes, *Rh3, Rh5, gbeta76, norpa, ninaC, ninaA, INAD, Calx, trpl, Arr1, Arr2* and *stops* (further discussed below; Figure 2E). Of the remaining eight annotation clusters, clusters 9 and 10 are also directly associated with vision and are enriched for homeobox and rhabdomere development, respectively. Two genes in common between these two clusters include *PvuII-PstI homology 13* (*Pph13*) and *ocelliless* (*oc*) that function in ocellus and compound eye photoreceptor development (Fichelson et al. 2012; Mahato et al. 2014).

Some of the genes enriched in other annotation clusters also have a role in vision. One gene in common between annotation clusters 4, 5 and 6 is *ora transientless* (*ort*), a gene that is necessary for vision as it encodes for a postsynaptic chlorine channel gated by the photoreceptor neutrotransmitter, histamine (Gengs et al. 2002). Annotation clusters 4, 5 and 8 include *resistant to dieldrin* (*Rdl*), a gene that has a role in the circuits underlying visual processing, odor coding, learning and memory, sleep and courtship behavior (Brotz et al. 2001; Liu et al. 2007; Chung et al. 2009; Yuan et al. 2013).

### Conservation of phototransduction genes in Lepidoptera

Genes commonly upregulated in heads were annotated with functions relating to vision and phototransduction (Figure 2E). Yet their evolutionary history and potential functional conservation requires further validation. To evaluate whether phototransduction genes were lost or expanded in Lepidoptera relative to *D. melanogaster*, we generated 32 insect phylogenies for 76 phototransduction-related genes including (Table S2-4). For each phylogeny, we searched 8 insect genomes including 2 moth species (*M. sexta* and *B. mori*) and 2 butterfly species (*H. melpomene* and *D. plexippus*). We predicted that we would find variation in phototransduction gene gains and loses between *D. melanogaster* and Lepidoptera and between moths and butterflies due to differences in eye morphology. Each *D. melanogaster* ommatidium consists of 8 photoreceptors with an open rhabdom, the structure where light is absorbed by the rhodopsins (Wernet et al. 2015). Unlike *D. melanogaster*, butterflies have 9 photoreceptor cells and a fused rhabdom (Wernet et al. 2015). Interestingly, moths and butterflies also differ from each other in eye morphology related to their light environments. Most butterflies have apposition-type eyes, where light from each lens is processed by one rhabdom and each ommatidium is separated by a sheath made of light-absorbing screening pigment to avoid light from other ommatidia (Kinoshita et al., 2017; Warrant & Dacke, 2016; Yack et al., 2007). Conversely, moths have superposition-type eyes where rhabdoms are separated from the crystalline cones by a translucent area allowing light to reach each rhabdom from hundreds of lenses (Warrant & Dacke, 2016; Yack et al., 2007).

Across all eight insect genomes we detect gene gains and losses in gene families such as opsin, trp, innexin and wunen (Figure 3A). Between *D. melanogaster* and lepidopterans, differences in gene gain and loss occur in the gene families opsin, innexin, wunen and DAGL. We did not detect any conserved differences in gene gain or loss between moths and butterflies (Figure 3A). Yet, an interesting gene family to note is *Vha100*, which has a *Vha100-like* gene that is lost in non-lepidopteran insects (Supplementary Results; Figure S5G) and also *innexin 9*, which is duplicated in *H. melpomene* (Supplementary Results; Figure S6). Since many of genes seem to be conserved between *D. melanogaster* and Lepidoptera, we visualized their expression in *M. sexta* heads and *H. melpomene* heads, antennae, legs and mouth parts. Upregulation of orthologs in *H. melpomene* heads would suggest a conserved role in vision for genes annotated with phototransduction function. Conversely, upregulation of a paralog suggests that butterflies are using a different member of the gene family to perform a visual function. We found 28 genes upregulated in heads relative to other tissue types (Figure 3B; Table 2; Figure S2-6). Most of the main genes involved in *D. melanogaster* phototransduction were found as single copies in Lepidoptera and were upregulated in *H. melpomene* heads such as *Gα, β* and *γ, norpA, inaD, ninaC, Calx, trp, trpl, Arr1, Arr2* and *stops* (Figure 1, Table 2). These results suggest that second messengers, ion channels, and termination of phototransduction are conserved between *D. melanogaster* and Lepidoptera (See below). The main differences in the phototransduction cascade between *H. melpomene* and *D. melanogaster* are in the opsins which initiate phototransduction and in DAG regulation (discussed further below; Figure 1). While there is no consistent difference between moths and butterflies in gene gains and losses, we found large differences in *trp* gene expression (See below).

**Table 2.**
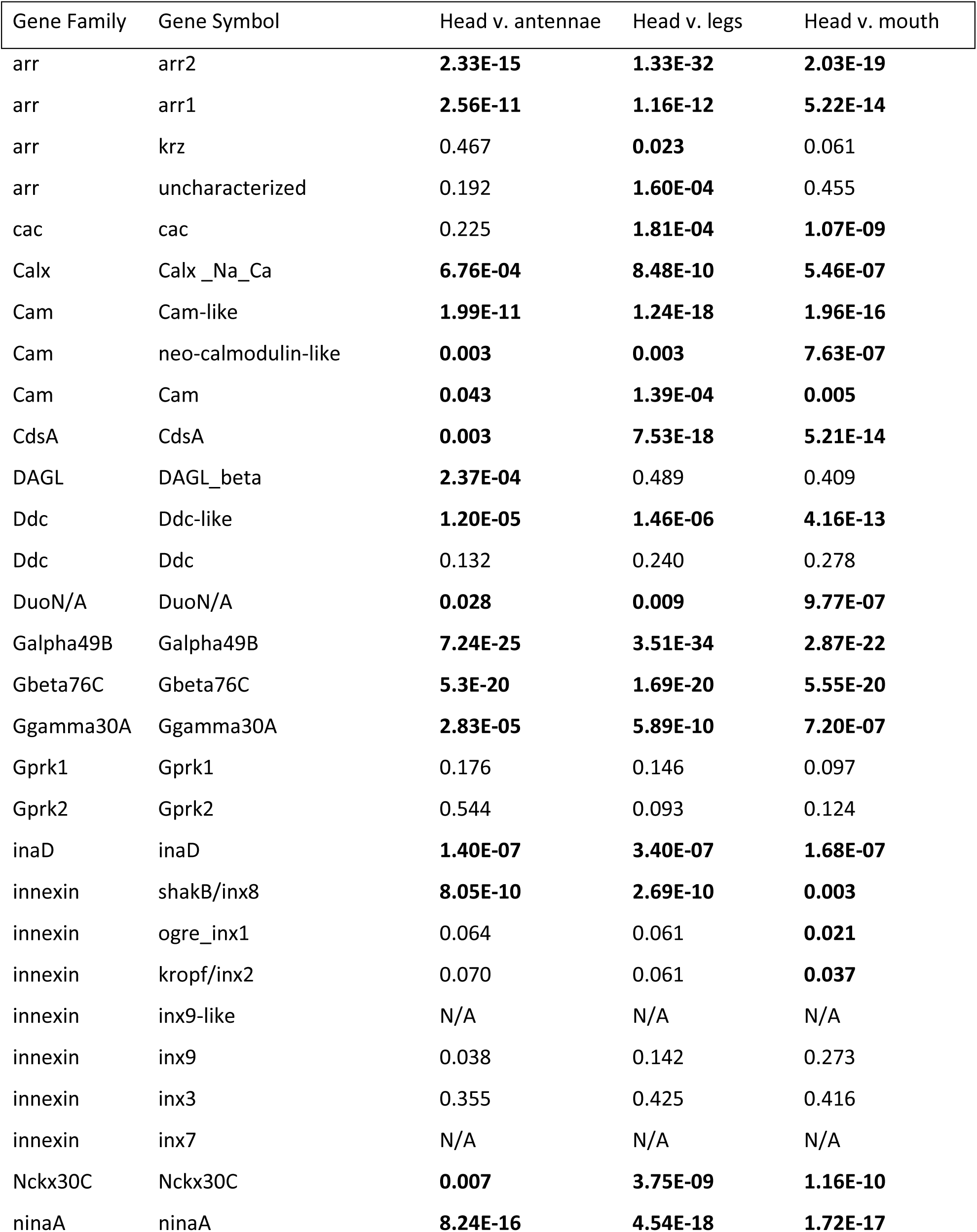
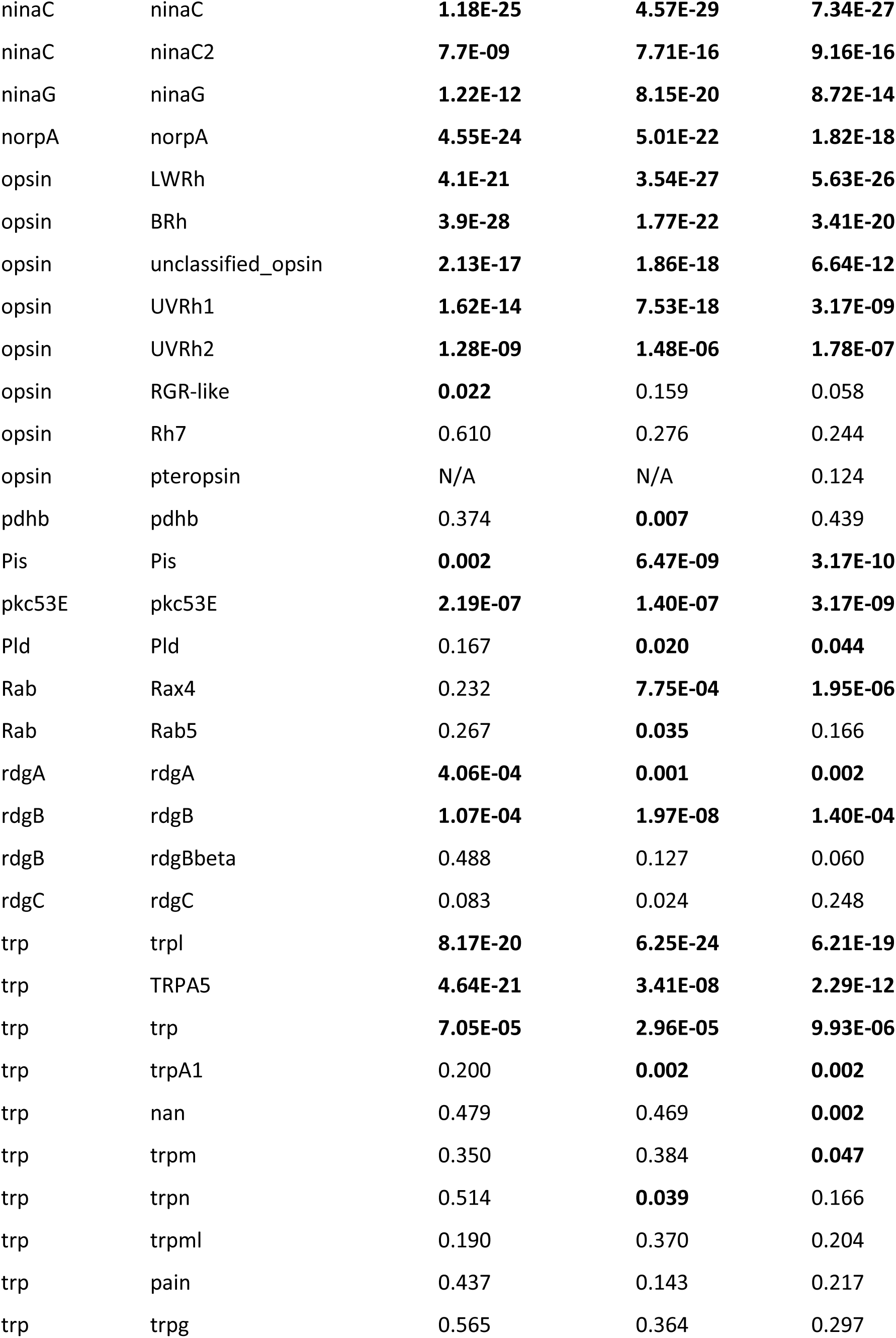
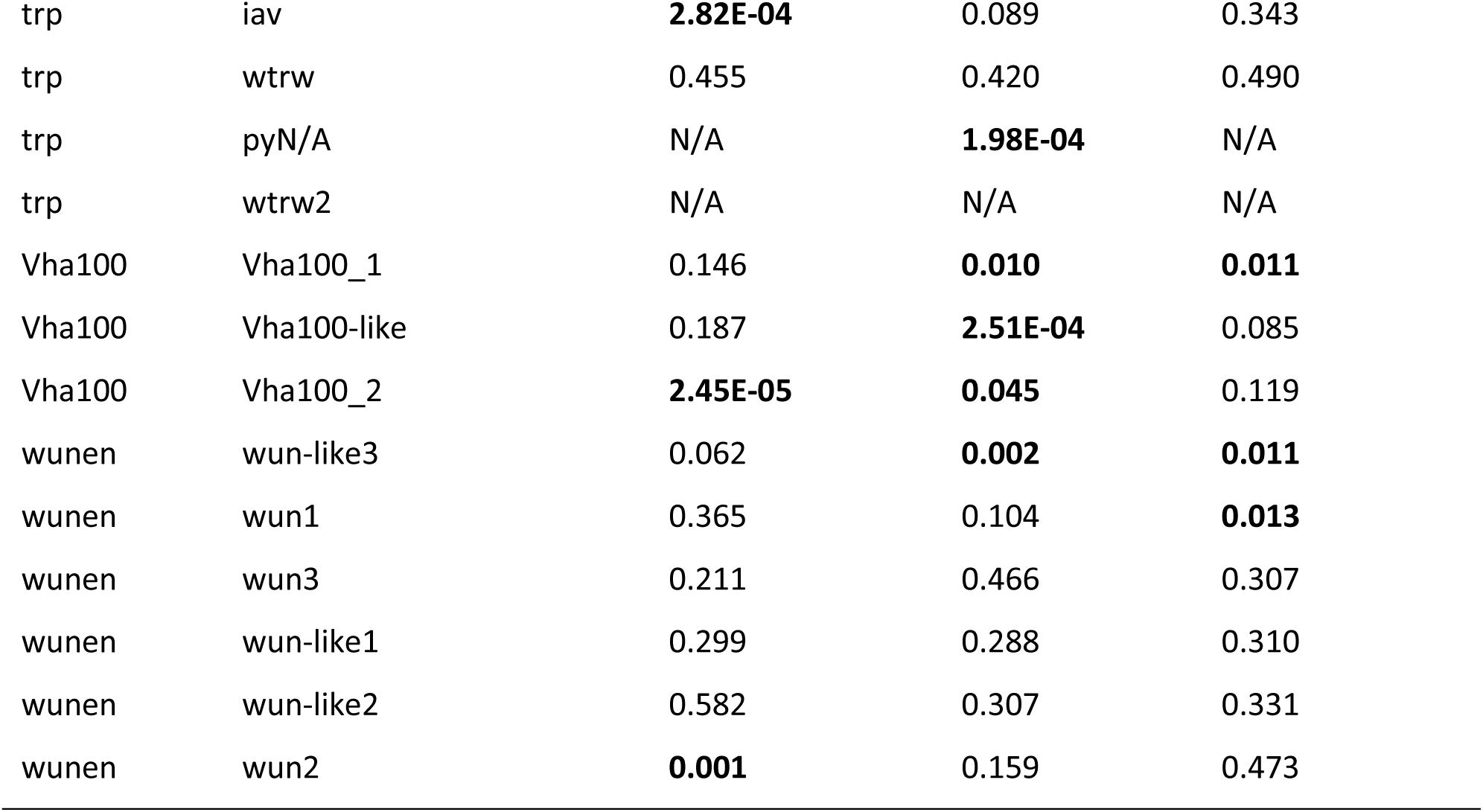
Q-values for four tissue pair-wise comparisons in *Heliconius melpomene*

| Gene Family | Gene Symbol | Head v. antennae | Head v. legs | Head v. mouth |
| --- | --- | --- | --- | --- |
| arr | arr2 | **2.33E-15** | **1.33E-32** | **2.03E-19** |
| arr | arr1 | **2.56E-11** | **1.16E-12** | **5.22E-14** |
| arr | krz | 0.467 | **0.023** | 0.061 |
| arr | uncharacterized | 0.192 | **1.60E-04** | 0.455 |
| cac | cac | 0.225 | **1.81E-04** | **1.07E-09** |
| Calx | Calx_Na_Ca | **6.76E-04** | **8.48E-10** | **5.46E-07** |
| Cam | Cam-like | **1.99E-11** | **1.24E-18** | **1.96E-16** |
| Cam | neo-calmodulin-like | **0.003** | **0.003** | **7.63E-07** |
| Cam | Cam | **0.043** | **1.39E-04** | **0.005** |
| CdsA | CdsA | **0.003** | **7.53E-18** | **5.21E-14** |
| DAGL | DAGL_beta | **2.37E-04** | 0.489 | 0.409 |
| Ddc | Ddc-like | **1.20E-05** | **1.46E-06** | **4.16E-13** |
| Ddc | Ddc | 0.132 | 0.240 | 0.278 |
| DuoN/A | DuoN/A | **0.028** | **0.009** | **9.77E-07** |
| Galpha49B | Galpha49B | **7.24E-25** | **3.51E-34** | **2.87E-22** |
| Gbeta76C | Gbeta76C | **5.3E-20** | **1.69E-20** | **5.55E-20** |
| Ggamma30A | Ggamma30A | **2.83E-05** | **5.89E-10** | **7.20E-07** |
| Gprk1 | Gprk1 | 0.176 | 0.146 | 0.097 |
| Gprk2 | Gprk2 | 0.544 | 0.093 | 0.124 |
| inaD | inaD | **1.40E-07** | **3.40E-07** | **1.68E-07** |
| innexin | shakB/inx8 | **8.05E-10** | **2.69E-10** | **0.003** |
| innexin | ogre_inx1 | 0.064 | 0.061 | **0.021** |
| innexin | kropf/inx2 | 0.070 | 0.061 | **0.037** |
| innexin | inx9-like | N/A | N/A | N/A |
| innexin | inx9 | 0.038 | 0.142 | 0.273 |
| innexin | inx3 | 0.355 | 0.425 | 0.416 |
| innexin | inx7 | N/A | N/A | N/A |
| Nckx30C | Nckx30C | **0.007** | **3.75E-09** | **1.16E-10** |
| ninaA | ninaA | **8.24E-16** | **4.54E-18** | **1.72E-17** |
| ninaC | ninaC | **1.18E-25** | **4.57E-29** | **7.34E-27** |
| ninaC | ninaC2 | **7.7E-09** | **7.71E-16** | **9.16E-16** |
| ninaG | ninaG | **1.22E-12** | **8.15E-20** | **8.72E-14** |
| norpA | norpA | **4.55E-24** | **5.01E-22** | **1.82E-18** |
| opsin | LWRh | **4.1E-21** | **3.54E-27** | **5.63E-26** |
| opsin | BRh | **3.9E-28** | **1.77E-22** | **3.41E-20** |
| opsin | unclassified_opsin | **2.13E-17** | **1.86E-18** | **6.64E-12** |
| opsin | UVRh1 | **1.62E-14** | **7.53E-18** | **3.17E-09** |
| opsin | UVRh2 | **1.28E-09** | **1.48E-06** | **1.78E-07** |
| opsin | RGR-like | **0.022** | 0.159 | 0.058 |
| opsin | Rh7 | 0.610 | 0.276 | 0.244 |
| opsin | pteropsin | N/A | N/A | 0.124 |
| pdhb | pdhb | 0.374 | **0.007** | 0.439 |
| Pis | Pis | **0.002** | **6.47E-09** | **3.17E-10** |
| pkc53E | pkc53E | **2.19E-07** | **1.40E-07** | **3.17E-09** |
| Pld | Pld | 0.167 | **0.020** | **0.044** |
| Rab | Rax4 | 0.232 | **7.75E-04** | **1.95E-06** |
| Rab | Rab5 | 0.267 | **0.035** | 0.166 |
| rdgA | rdgA | **4.06E-04** | **0.001** | **0.002** |
| rdgB | rdgB | **1.07E-04** | **1.97E-08** | **1.40E-04** |
| rdgB | rdgBbeta | 0.488 | 0.127 | 0.060 |
| rdgC | rdgC | 0.083 | 0.024 | 0.248 |
| trp | trpl | **8.17E-20** | **6.25E-24** | **6.21E-19** |
| trp | TRPA5 | **4.64E-21** | **3.41E-08** | **2.29E-12** |
| trp | trp | **7.05E-05** | **2.96E-05** | **9.93E-06** |
| trp | trpA1 | 0.200 | **0.002** | **0.002** |
| trp | nan | 0.479 | 0.469 | **0.002** |
| trp | trpm | 0.350 | 0.384 | **0.047** |
| trp | trpn | 0.514 | **0.039** | 0.166 |
| trp | trpml | 0.190 | 0.370 | 0.204 |
| trp | pain | 0.437 | 0.143 | 0.217 |
| trp | trpg | 0.565 | 0.364 | 0.297 |
| trp | iav | **2.82E-04** | 0.089 | 0.343 |
| trp | wtrw | 0.455 | 0.420 | 0.490 |
| trp | pyN/A | N/A | **1.98E-04** | N/A |
| trp | wtrw2 | N/A | N/A | N/A |
| Vha100 | Vha100_1 | 0.146 | **0.010** | **0.011** |
| Vha100 | Vha100-like | 0.187 | **2.51E-04** | 0.085 |
| Vha100 | Vha100_2 | **2.45E-05** | **0.045** | 0.119 |
| wunen | wun-like3 | 0.062 | **0.002** | **0.011** |
| wunen | wun1 | 0.365 | 0.104 | **0.013** |
| wunen | wun3 | 0.211 | 0.466 | 0.307 |
| wunen | wun-like1 | 0.299 | 0.288 | 0.310 |
| wunen | wun-like2 | 0.582 | 0.307 | 0.331 |
| wunen | wun2 | **0.001** | 0.159 | 0.473 |

**Figure 3:**
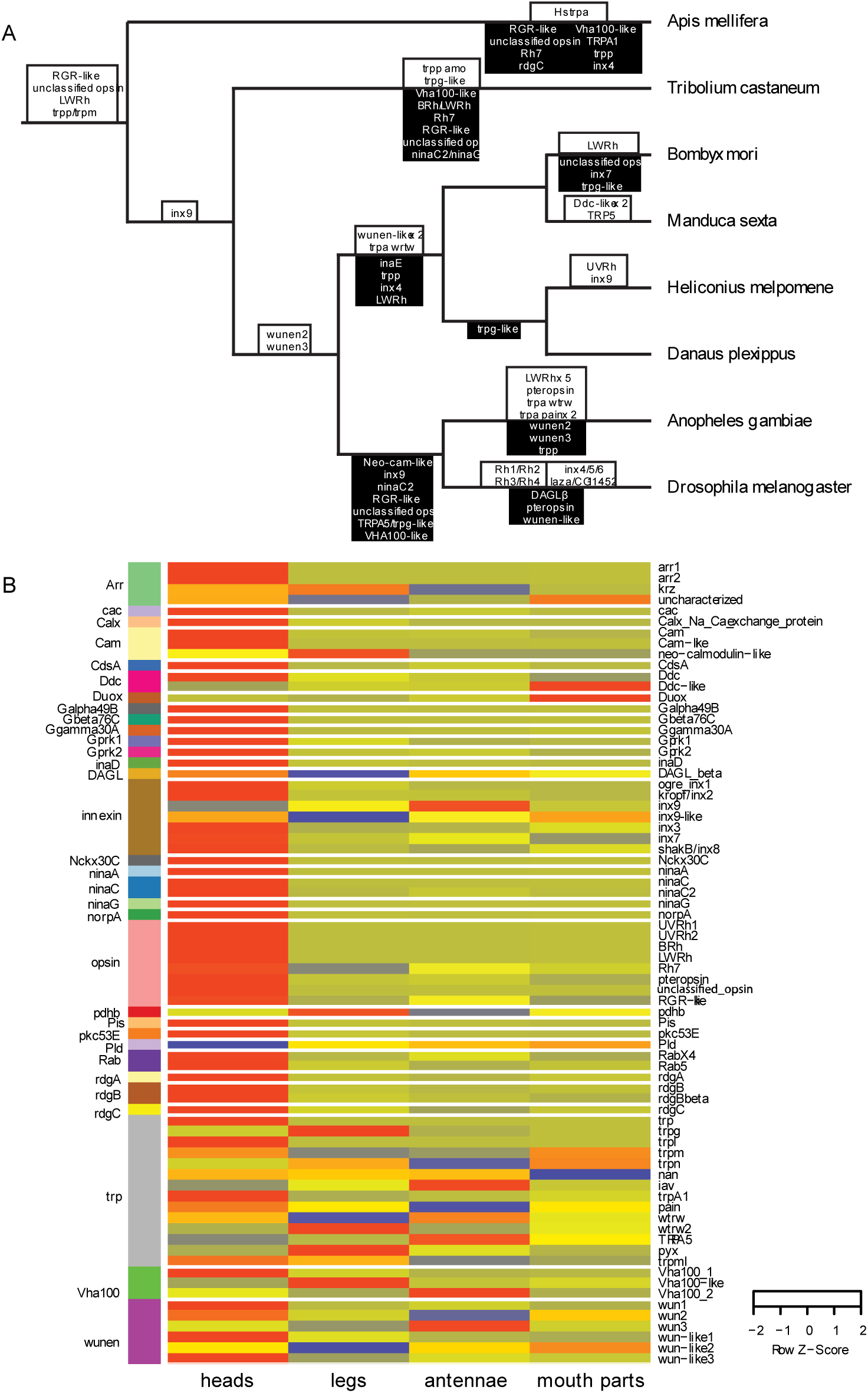
Phototransduction genes gains, losses, and expression. (A) Insect phylogeny showing gains in white boxes above tree branch and losses in black boxes below tree branch. (B) Heatmap of expression of genes orthologous to *D. melanogaster* phototransduction genes in tissues of *H. melpomene* heads, antennae, legs and mouth parts. Red signifies higher expression while blue signifies downregulation. Gene names are listed on the right while gene family names are listed on the left and assigned a different block color per gene family.

### Opsins in Lepidoptera

We began our survey of phototransduction genes in Lepidoptera by investigating the molecular evolution and expression of opsin genes typically responsible for initiating the phototransduction cascade (Figure 1). Opsin phylogenies have been the focus of many studies attempting to understand the evolutionary history of light detection (Arendt 2003; Raible et al. 2006; Plachetzki et al. 2007; Suga et al. 2008; Porter et al. 2012; Ramirez et al. 2016; Vöcking et al. 2017). These studies have reconstructed opsin presence in the ancestor of bilaterian animals (Ramirez et al. 2016) and have described a new opsin type (Vöcking et al. 2017). To inspect the phylogenetic history of the opsins, we added *H. melpomene* sequences from the reference genome and a *de novo* transcriptome to a set of sequences used in Kanost et al. (2016). We recovered the previously described *Heliconius-*specific *UVRh* duplication and orthologs for all other known opsins (Figure 4A) (Briscoe et al. 2010; Yuan et al. 2010; McCulloch et al. 2017). We also found two opsin genes: an *unclassified opsin (UnRh)* first described in Kanost et al. (2016) and *RGR-like* that both lacked a *D. melanogaster* ortholog but were found in our butterfly genomes (Figure 4A).

**Figure 4:**
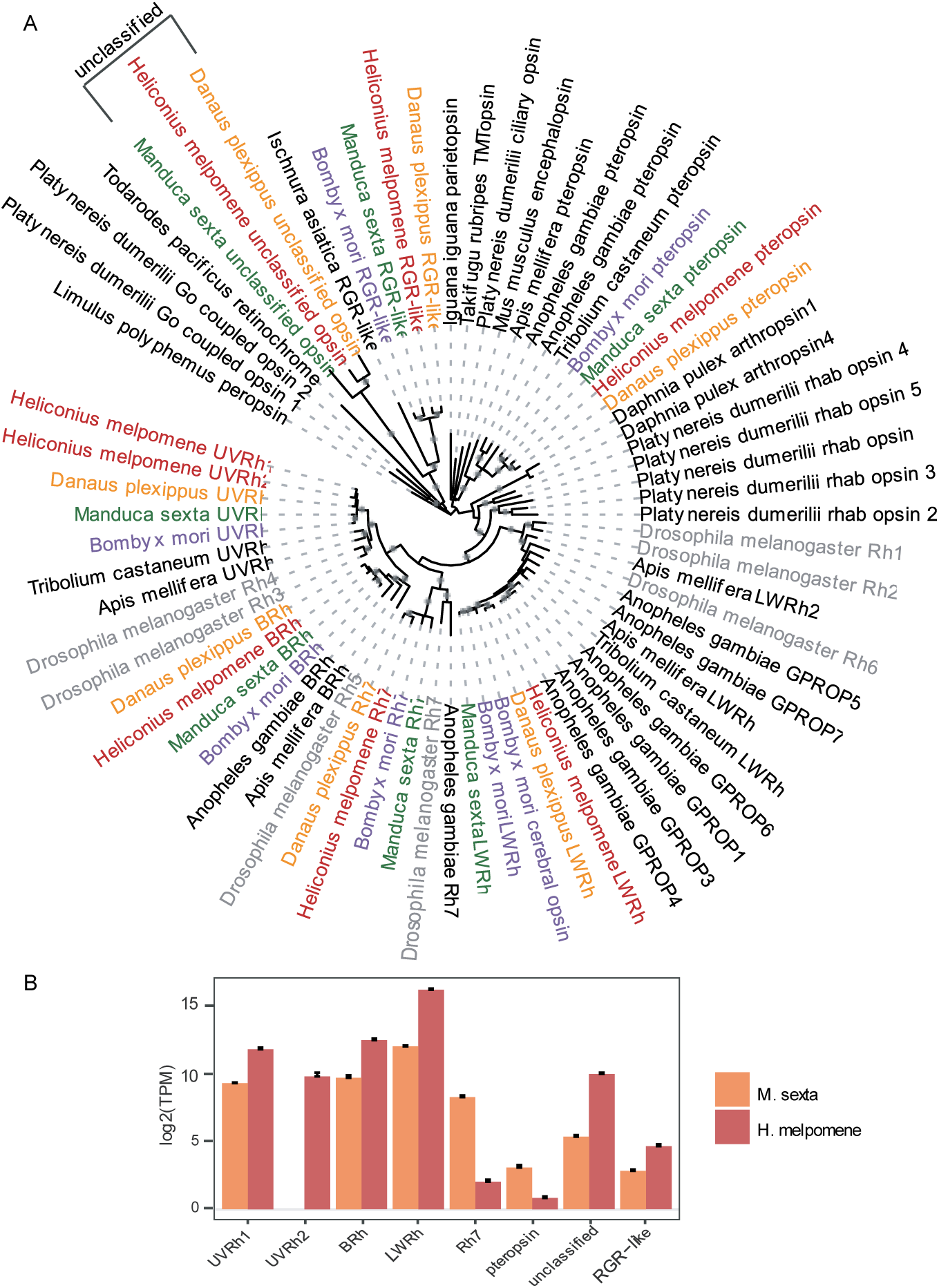
Insect opsin phylogeny and opsin gene expression in a moth and butterfly. (A) Opsin phylogenetic tree generated using sequences from Kanost et al. (2016) and sequences from *H. melpomene* and *D. plexippus*. (B) Expression of opsin genes in *M. sexta* heads (n=8) and *H. melpomene* heads (n=8).

To predict the role of all opsin genes we looked at their expression profile in *M. sexta* and *H. melpomene.* In *M. sexta*, all opsins had expression in head tissue (Figure 4B). In *H. melpomene*, our functional enrichment showed that homologs of *D. melanogaster* rhodopsin genes *Rhodopsin 3* (*Rh3*) and *Rhodopsin 5* (*Rh5*), which correspond to *UVRh1/Rh2* and *BRh* respectively were upregulated in *H. melpomene* heads (Figure 2E; Table S11) (Briscoe et al. 2010; Yuan et al. 2010). *LWRh* and the *unclassified opsin* are also upregulated in *H. melpomene* heads (Table 2; Figure 4B). *LWRh* was the most highly expressed opsin gene probably due to the amount of LW photoreceptor cells per ommatidium. *Heliconius* have 9 photoreceptor cells where at least six cells express *LWRh* and two express short wavelength *BRh, UVRh1* or *UVRh2* (McCulloch et al. 2016, 2017).

Upregulation of the *unclassified opsin* (*UnRh*) was intriguing because Kanost et al (2016) noted the unclassified opsin lacks a lysine at the typical location where the chromophore is bound in opsins, yet the gene is highly expressed in *H. melpomene* eyes and brain suggesting a role in vision. A recent study found that alternative amino acids sites may be used in some G-protein coupled receptors for chromophore-binding (Faggionato & Serb 2017). Further, cephalopods have a photosensitive pigment called retinochrome, studied biochemically, that lacks a conserved rhodopsin glutamic acid base (Terakita et al. 1989, 2000). Retinochrome, unlike rhodopsin, binds an all-*trans* retinal and acts as a photoisomerase converting the chromophore to 11-*cis* to regenerate the photosensitive rhodopsin (Sperling & Hubbard 1975). Not only are rhodopsin and retinochrome important for the cephalopod visual system, they are also found together in extraocular tissues serving a potential role in camouflage (Kingston et al. 2015). By adding a squid retinochrome sequence to our opsin phylogeny we found that the lepidopteran-specific unclassified opsin and RGR-like opsin are more closely-related to retinochrome than they are to other opsins with known functions (Figure 4A).

Notably, Lepidoptera are thought to rely more on enzymatic regeneration of 11-*cis*-retinal than is the case in Diptera (Bernard 1983a and b; Stavenga and Hardie 2011). As the unclassified opsin has high expression in eyes and is phylogenetically similar to retinochrome, both proteins may have related enzymatic roles in vision if the unclassified opsin is expressed near the photoreceptor cells. To localize where in the butterfly eye the unclassified opsin is expressed, we made an antibody against the unclassified opsin. We visualized unclassified opsin expression relative to that of LWRh opsin in *H. melpomene*. In *Heliconius*, LWRh is expressed in photoreceptor cells R3-8 (McCulloch et al. 2016). Intriguingly, we found the unclassified opsin protein product abundantly expressed in crystaline cone cells, in primary pigment cells, and in the six secondary pigment cells surrounding the ommatidium (Figure 5). Staining is brighter in the distal part of the retina presumably because the secondary pigment cells decrease in size as they approach the basement membrane (Figure 5A). If the unclassified opsin had a function similar to that of the color vision opsins, we would expect it to be expressed in the photoreceptor cells. However, this protein is expressed in other retinal cells adjacent to the photoreceptor cells.

**Figure 5:**
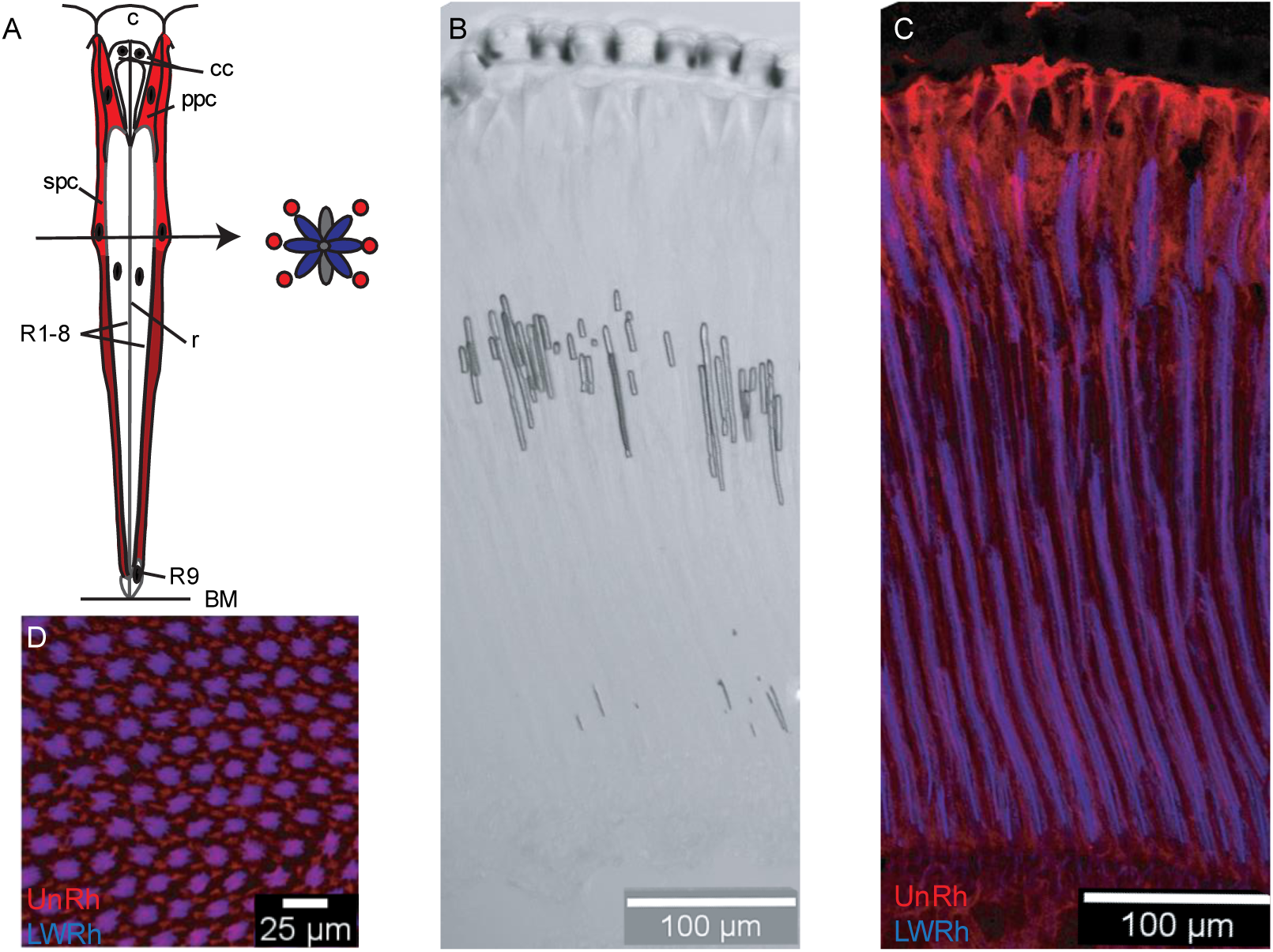
Immunohistochemistry of a butterfly retinochrome, unclassified opsin (UnRh). (A) Drawing of a butterfly ommatidium showing the cornea (c), crystalline cone (cc), rhabdom (r), photoreceptor cells (R1-9), primary pigment cells (ppc), secondary pigment cells (spc), basal pigments cells (bpc) and basement membrane (bm) based on Kolb (1985). Red represents areas where UnRh expression is detected, dark red indicates where the cell presumably narrows and staining is not as bright. A drawing of a cross section shows cells R1-8, blue cells represent staining of LWRh and red circles represent UnRh expression. (B) Brightfield image of a longitudinal section of a butterfly eye showing the anatomy of each ommatidium. (C) Longitudinal section of a *H. melpomene* retina stained for opsins LWRh (blue) and UnRh (red). (D) Transverse section stained for opsins LWRh (blue) and UnRh (red).

In squid, the retinochrome is expressed in inner segment cells while the rhodopsin that it interchanges chromophore with is in the outer segment, separated by the basement membrane (Kingston et al. 2015; Chung & Marshall 2017). Taken together, these results suggest that the unclassified opsin might have a role similar to the squid retinochrome in photoisomerization of the butterfly chromophore. This mechanism could be required for fast regeneration of an active rhodopsin necessary to quickly process color information during flight.

### Regulation of diacylglycerol (DAG)

After phototransduction is triggered by photon absorption, G_q_*α* is released from a G-protein complex of 3 subunits (*α, β*, and *γ*) and activates phospholipase C (PLC encoded by *norpA*) to produces diacylglycerol (DAG) (Bloomquist et al. 1988; Lee et al. 1994). DAG has been implicated in the activation of transient receptor potential (TRP and TRPL) channels (Leung et al. 2008; Chyb et al. 1999). DAG is hydrolyzed is by the actions of DAG lipase (DAGL) encoded by the gene *inaE* (Leung et al. 2008). *InaE* mutants in *D. melanogaster* have defective responses to light, demonstrating that DAGL activity is required for photoreceptor responses (Leung et al. 2008). Although this gene is crucial for *D. melanogaster* phototransduction, an ortholog of *inaE* is missing in Lepidoptera (Figure 3A). We found that Lepidoptera retain *DAGLβ, D. melanogaster* retains *DAGLα* (*inaE*), and *A. mellifera, A. gambiae, T. castaneum*, and mammals retain both (Figure 6A). Both *DAGLα* and *DAGLβ* encode an Sn-1 diacylglycerol lipase that generate a monoacylglycerol (MAG) product. Note that for *T. castaneum, DAGLα* is not included in the phylogeny because the sequence was too short to generate a correct alignment. We predict that *DAGLβ* carries out the phototransduction function of hydrolyzing DAG in moth and butterfly vision because Lepidoptera have lost an ortholog of *D. melanogaster inaE* and retained *DAGLβ*. *DALGβ* was expressed in *M. sexta* heads and in *H. melpomene* heads (Figure 6B). While we confirm expression in heads, *DAGLβ* is not upregulated in heads relative to other tissue types. *DAGLβ* may have a role in vision in Lepidoptera, but it might also be used in other tissues for other functions.

**Figure 6:**
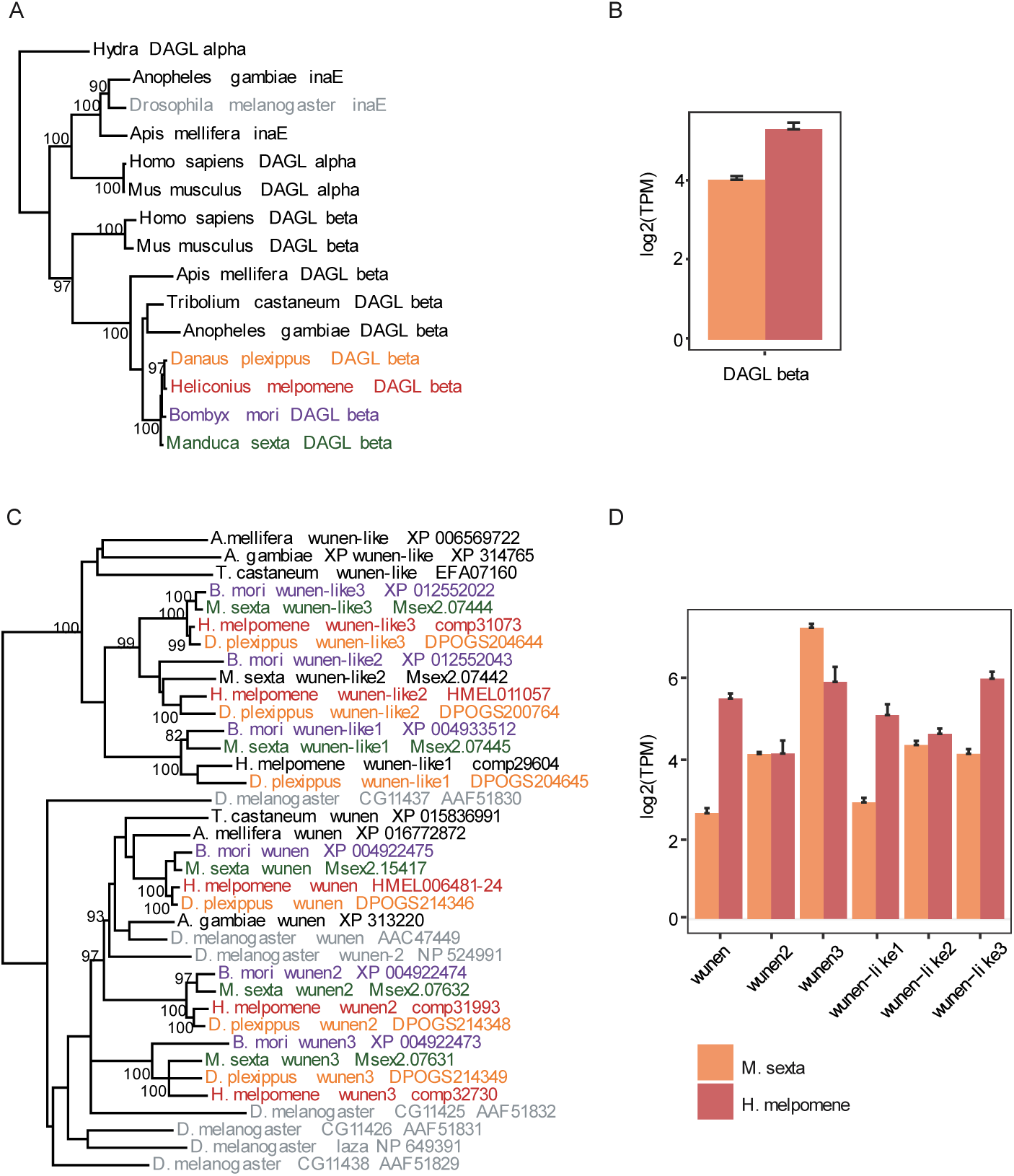
Molecular evolution and expression of DAGL and wunen. (A) *DAGL* phylogenetic tree generated using amino acid sequences from 8 insect genomes and *H. sapien* and *M. musculus.* (B) Expression of DAGL genes in *M. sexta* heads (n=8) and *H. melpomene* heads (n=8). (C) *Wunen* phylogenetic tree generated using sequences from 8 insect genomes. (D) Expression of wunen genes in *M. sexta* heads (n=8) and *H. melpomene* heads (n=8).

DAG level is also regulated by degeneration A (RDGA) (conserved in moths and butterflies, Figure S5) and Lazaro (LAZA) (Garcia-Murillas et al. 2006; Bao & Friedrich 2009). Lazaro is a lipid phosphate phosphatase (LPP) and is found in *D. melanogaster* photoreceptors (Garcia-Murillas et al. 2006). *Lazaro* is a member of the *wunen* subfamily (Figure 6C). *Wunen* helps regulate the level of bioactive phospholipids, has a role in germ line migration and is necessary for tracheal development (Ile et al. 2012; Zhang et al. 1997). We found 7 sequences belonging to the *wunen* gene family in *D. melanogaster*; *Lazaro* is a *D. melanogaster-*specific duplication. While other non-*D. melanogaster* insects have one copy of *wunen*, lepidopterans had 3 copies (Figure 6C). In addition, while other insects had one copy of *wunen-like*, Lepidoptera had 3 copies of *wunen-like* that arose after lepidopteran divergence from other insects (Figure 6C). All copies of *wunen* and *wunen-like* were expressed in *M. sexta* and *H. melpomene* heads (Figure 6C). *Wunen* and *wunen-like-3* were the two copies most highly expressed in *H. melpomene* heads.

### TRP and TRPL

Transient receptor potential (TRP and TRPL) channels are essential in *D. melanogaster* phototransduction. They allow the influx of Ca^2+^ and cause cell depolarization (Montell & Rubin 1989). *Trp* is the dominant light-sensitive channel in the rhabdomeres (∼10x more abundant than *trpl*), and flies with mutated *trp* behave as though they are blind (Montell & Rubin 1989). The TRP superfamily contains more than 20 cation channels (Montell et al. 2002). While *trp* and *trpl* function in *D. melanogaster* vision, other *trp* genes sense pain, vanilloid compounds, and heat, among other stimuli (Montell et al. 2002; Montell 2005). In our examination of the TRP gene family, we found 14 genes members of this gene family in *H. melpomene* and 17 in *M. sexta*.

Differences between *D. melanogaster* and Lepidoptera included a duplication of *trpa wtrw* and loss of *trpp* in moths and butterflies. The function of *trpa wtrw* (encoding TRP channel water witch) has not been characterized in any insect species but *trpa* genes, related gene family members, have been shown to function in temperature sensitivity, fructose aversion, and sexual receptivity in *D. melanogaster* (Xu et al. 2008; Sakai et al. 2009; Peng et al. 2016). *Trpa wtrw* is expressed in *M. sexta* heads and in *H. melpomene* heads whereas *trpa wtrw2* has very low expression. In the *trp* family, *M. sexta* retain a *trpg-like* gene that is lost in *D. melanogaster* and butterflies *H. melpomene* and *D. plexippus* (Figure 3A; Figure 7). *Trpg* encodes a protein that is found in *D. melanogaster* photoreceptors and has been speculated to form a heteromultimeric channel with TRPL (Montell 2005). *Trpg* role in *Drosophila* vision is uncertain, although it is expressed in *H. melpomene* heads; both *trpg* and *trpg-like* have low expression in *M. sexta* heads (Figure 7B). Furthermore, *M. sexta* also has 3 copies of *TRPA5* which is lost in *D. melanogaster* and *A. gambiae* (Figure 7A). *TRPA5* genes in humans and mice are predominantly expressed in the brain, and it has been suggested that they might function in learning and memory (Okada et al. 1998; Fowler et al. 2007). All three copies of *TRPA5* are expressed *M. sexta* heads, and *TRPA5* is also expressed in *H. melpomene* heads (Figure 7B).

**Figure 7:**
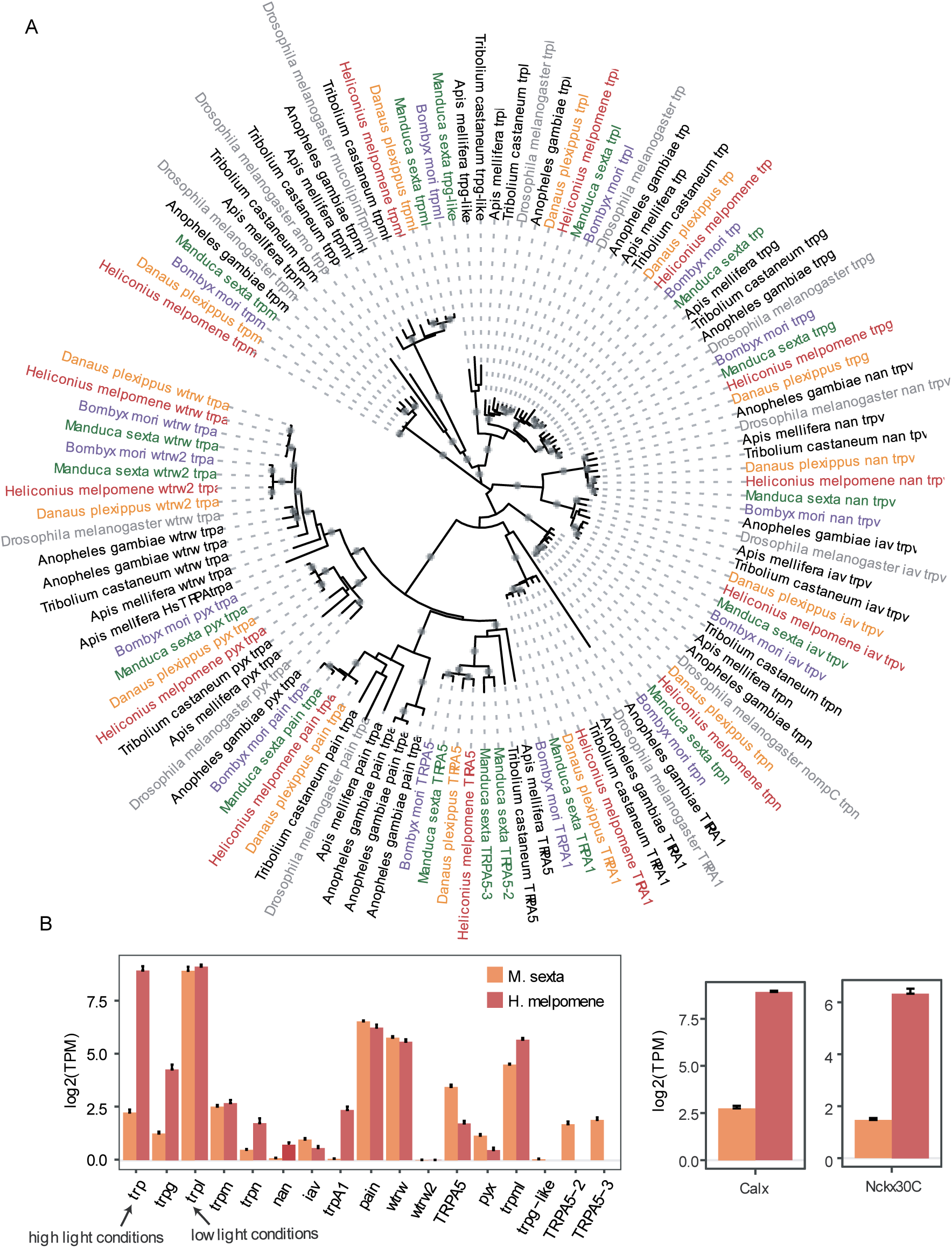
Molecular evolution and expression of the transient receptor potential (trp) cation channel gene family. (A) *Trp* phylogenetic tree generated using sequences from 8 insect genomes. (B) Expression of *trp, Calx*, and *Nckx30C* genes in *M. sexta* heads (n=8) and *H. melpomene* heads (n=8). TRP and TRPL used in high light and low light conditions respectively are highlighted.

### Ion channels used in diurnal and nocturnal insects

A transcriptome study in cockroaches found that *trpl* was approximately 10 times more abundant than *trp* (French et al. 2015). RNAi of *trpl* reduced electroretinogram (ERG) response much more than *trp* after 21 days suggesting that, unlike *D. melanogaster*, cockroach TRPL rather than TRP has a larger contribution to phototransduction (French et al. 2015). The authors suggested that differences in visual ecology are responsible for differential functions of the ion channels: daylight-active *D. melanogaster* rely on fast responsive trp and dark- or dim-light active cockroaches rely on trpl (French et al. 2015). We found that *trp* and *trpl* are both highly expressed in *H. melpomene* heads which is different from either *D. melanogaster* or cockroach. Like cockroaches, we found that *trp* and *trpl* both have expression in *M. sexta* heads, but *trp* is expressed at a much lower level compared to *trpl* (Figure 7B). Our results suggest that the TRPL ion channel is also used by Lepidoptera in low light conditions.

TRP and TRPL channels that allow Ca^2+^ and Na^+^ into the photoreceptor cell are co-localized with an Na^+^/Ca^2+^ exchanger encoded by *Calx* which allow Ca^2+^ out of the cell (Montell 2005). Mutations of *Calx* result in a transient light response and a decrease in signal amplification implying a role for this gene in Ca^2+^ maintenance for proper TRP signaling (Wang et al. 2005). Overexpression of *Calx* can suppress retinal degeneration due to TRP constitutive activation (Montell 2005; Wang et al. 2005). *Calx* is upregulated in *H. melpomene* heads and found as a single copy in all insect genomes (Figure 2E; Table 2; Figure 7; Figure S2C). We detected a lower expression of *Calx* in *M. sexta* heads compared to the expression in *H. melpomene* heads potentially correlated with the lower expression of *trp* compared to *trpl* in *M. sexta* (Figure S2C). A similar pattern of expression was also observed for another Na^+^/Ca^2+^ exchanger encoded by *Nckx30C*. *Nckx30C* was upregulated in *H. melpomene* heads yet expression of this ion channel was lower in *M. sexta* heads compared to *H. melpomene* heads (Table 2; Figure 7; Figure S4B). *Nckx30C* has a similar role to *Calx* in moving Ca^2+^ out of the cell (Haug-Collet et al. 1999). Both *Nckx30C* and *Calx* are expressed in the embryonic nervous system of *D. melanogaster* and in the adult eye and brain (Haug-Collet et al. 1999). Our results suggest that decreased expression of *trp* in nocturnal moths is accompanied by a decrease in *Calx* and *Nckx30C*. We conclude that one difference between moth and butterfly phototransduction appears to be in the expression of ion channels used for calcium exchange.

### Proposed Phototransduction Cascade in Lepidoptera

Based on phylogenetic relationships and gene expression we propose a model of phototransduction in Lepidoptera (Figure 1). Phototransduction initiation requires an opsin to be bound to a chromophore to initiate the cascade. We propose that in Lepidoptera, the chromophore is transported by CTD31 rather than the ortholog of *D. melanogaster* PINTA, which has been lost in lepidopterans (Macias-Muñoz et al. 2017). Similar to *D. melanogaster*, visual opsins (*BRh, LWRh*, and *UVRh*) initiate the phototransduction cascade by a change in configuration when the chromophore molecule absorbs light energy (von Lintig et al. 2010). We note that lepidopterans vary in opsin number (Frentiu et al. 2007; Briscoe 2008; Pirih et al. 2010; Xu et al. 2013). Photoisomerized 11-*cis*-3-hydroxyretinal is supplied to light-activated rhodopsin by retinochrome (UnRh) proteins found in pigment cells. Change in opsin conformation due to light absorption triggers the G-protein signaling cascade. Gqα, β, and γ are present as single copies and highly expressed in heads suggesting a conserved function in phospholipase C activation, encoded by *norpA*, when *G*_*q*_*α* is released (Figure 2E; Figure S3D-F; Figure S4E) (Bloomquist et al. 1988; Lee et al. 1994).

Phosopholipase C (PLC) produces inositol 1,4,5-trisphosphate (InsP_3_) and diacylglycerol (DAG) (Bloomquist et al. 1988; Hardie 2001). However, the regulation of DAG levels appears to differ between lepidopterans and *D. melanogaster* due to a loss of *laza* and *inaE*. We proposed that in Lepidoptera the actions of *inaE* are undertaken by a lepidopteran paralog *DAGL-beta* and those of *laza* by other members of the gene family, *wunen* or *wunen-like3*. LAZA acts in opposition to DAG kinase encoded by rdgA (Garcia-Murillas et al. 2006). In *D. melanogaster*, DAG is converted into PIP_2_ by the phosphoinositide pathway which gives photoreceptor cells sensitivity and fast response (Hardie 2001; Garcia-Murillas et al. 2006). We believe that the actions of this pathway remain conserved in Lepidoptera because *rdgA, cdsA*, and *rdgB* are upregulated in *H. melpomene* heads. While phosphatidic acid (PA) is likely converted into DAG by a *laza* paralog (*wunen* or *wunen-like3*), kinase *rgdA* maintains a role in converting DAG into PA. *CDP-diacylglycerol synthase* encodes a protein that converts PA into cytidine diphosphate diacylglycerol (CDP-DAG). Phosphatidyl inositol (PI) synthase then changes CDP-DAG into PI which is transported by phosphatidylinositol transfer protein encoded by (*rdgB*). Phosphorylation converts PI into PIP_2_. The actions by which DAG functions in phototransduction are not well understood. DAG lipase produces the metabolite polyunsaturated monoacylglycerol (MAG) (Montell 2012). It is thought that DAG might activate TRP and TRPL channels, although its role in phototransduction is debated (Leung et al. 2008; Chyb et al. 1999)

TRP and TRPL allow Ca^2+^ and Na^+^ into the cell which causes the photoreceptor cell to depolarize (Montell & Rubin 1989). We propose that the phototransduction cascade varies between moths and butterflies in the deployment of TRP and Na^+^/Ca^+^ channels. According to our expression data, butterflies use TRP and TRPL in similar amounts, while moths downregulate their TRP channel mRNAs. Since moths presumably have less TRP channels allowing in Ca^2+^, they also downregulate Na^+^/Ca^2+^ channels encoded by *Calx* and *Nckx30C*.

Phototransduction requires protein complexes to transduce and terminate the signal. One such complex is a target of G*αq* and is formed by INAD, TRP, PLC, and protein kinase C (PCK) (Shieh et al. 1989; Chevesich et al. 1997; Tsunoda et al. 1997; Bähner et al. 2000; Montell 2005). *InaD* is required to localize and coordinate proteins in the phototransduction cascade to the microvillar membrane (Bähner et al. 2000). The INAD and NinaC bind to each other, and individually bind calmodulin, which accelerates arrestin binding to rhodopsin to terminate phototransduction (Liu et al. 2008; Venkatachalam et al. 2010). Arrestin 1 and Arrestin 2 bind light-activated rhodopsin and discontinue cascade signaling in *D. melanogaster* (Stavenga & Hardie 2011; Dolph et al. 1993). Our data suggests that Arrestin 2 might be the major arrestin in butterfly phototransduction; it is more highly expressed than Arrestin 1 in moths as well, although to a lesser degree (Figure S2A). Phototransduction is also terminated by a protein with a suppressor of cytokine signaling (SOCS) box encoded by *stops*. The *stops* phenotype is associated with slow termination of phototransduction due to a decrease in NORPA (PLC) (Wang et al. 2008). We find these genes to be upregulated in butterfly heads (Figure 2B and Table 2), suggesting the actions of these complexes remain conserved. Lastly, Lepidoptera have a *ninaC2* gene, missing in *D. melanogaster*, which is upregulated in *H. melpomene* heads (Figure S4G).

## Conclusions

Most studies of phototransduction in insects extrapolate from what is known in *D. melanogaster* to assign potential functions to genes based on sequence similarity. In our study, we used transcriptomics and phylogenetics to explore the conservation of phototransduction genes between *D. melanogaster* and Lepidoptera. We found that many orthologs of key *D. melanogaster* phototransduction genes were upregulated in *H. melpomene* heads relative to legs, antennae, and mouth parts. Our results suggest that many features of *D. melanogaster* phototransduction cascade are conserved in lepidopteran vision. However, we found instances where lepidopteran paralogs are implicated in carrying out a role in vision when an ortholog is lost. Differences in phototransduction between *D. melanogaster* and Lepidoptera occur in chromophore transport, chromophore regeneration, opsins, and DAG regulation. While we found no conserved differences between moths and butterflies in gene gains and losses, quantifying gene expression in *M. sexta* and *H. melpomene* allowed us to detect differences in phototransduction between moth and butterflies. Notably, we found evidence that butterflies use both TRP and TRPL channels for phototransduction while moths downregulate trp, which is used for high light conditions (French et al. 2015). Along with decreased expression of *trp*, Na^+^/Ca^2+^ exchange proteins appear to show decreased expression in nocturnal moths. We have thus completed the first investigation of the evolution of the phototransduction cascade in Lepidoptera and found that differences between Lepidoptera and *D. melanogaster* are due to gene gain and loss while differences between moths and butterflies are due to gene expression changes.

## Acknowledgements

We thank Ali Mortazavi, Kevin Thornton, Jorge Llorente-Bousquets and Pablo Vinuesa for advice on data analysis. We thank Zachary Johnston, JP Lawrence, and Andrew Dang for comments on the manuscript, and Roger Hardie for extremely helpful discussions. This work was supported in part by National Science Foundation grants DEB-1342759 and IOS-1656260 to A.D.B., a Ford Foundation Predoctoral Fellowship to AMM, and a CNBES grant to AGRO.

---

**Figure S1: Heatmaps for four tissue pair-wise comparisons.** (A) Heatmap for differentially expressed (DE) genes between head and antennae. (B) Heatmap for DE genes between head and legs. (C) Heatmap for DE genes between head and mouth parts. DE genes were found using a Bonferroni correction.

**Figure S2: Phylogeny and expression of phototransduction genes 1.** Phylogenetic trees were generated using sequences from 8 insect genomes. Expression for all genes follows the convention boxed in red; orange bars represent expression in *M.sexta* heads, red bars expression *H. melpomene* heads, lilac expression in *H. melpomene* legs, green expression in *H. melpomene* antennae, and purple expression in *H. melpomene* mouth parts. (A) Arrestin gene family. (B) Cacophony gene family. (C) Calx Na^+^/Ca^+^ exchange protein gene family. (D) Calmodulin gene family.

**Figure S3: Phylogeny and expression of phototransduction genes 2.** Phylogenetic trees were generated using sequences from 8 insect genomes. Orange bars represent expression in *M. sexta* heads, red bars expression *H. melpomene* heads, lilac expression in *H. melpomene* legs, green expression in *H. melpomene* antennae, and purple expression in *H. melpomene* mouth parts. (A) CDP-diacylglycerol synthase gene family. (B) Dopa decarboxylase gene family. (C) Dual oxidase gene family. (D) G protein alpha q subunit gene family. (E) G protein beta subunit 76C gene family. (F) G protein subunit at 30A gene family. (G) G protein-coupled receptor kinase 1 gene family. (H) G protein-coupled receptor kinase 2 gene family.

**Figure S4: Phylogeny and expression of phototransduction genes 3.** Phylogenetic trees were generated using sequences from 8 insect genomes. Orange bars represent expression in *M. sexta* heads, red bars expression *H. melpomene* heads, lilac expression in *H. melpomene* legs, green expression in *H. melpomene* antennae, and purple expression in *H. melpomene* mouth parts. (A) Inactivation no afterpotential D gene family. (B) Nckx30C gene family. (C) Neither inactivation nor afterpotential A gene family. (D) NinaG gene family. (E) No receptor potential A gene family. (F) Pyruvate dehydrogenase E1 beta subunit gene family. (G) Neither inactivation nor afterpotential C gene family. (H) RabX4 and Rab5 gene family.

**Figure S5: Phylogeny and expression of phototransduction genes 4.** Phylogenetic trees were generated using sequences from 8 insect genomes. Orange bars represent expression in *M. sexta* heads, red bars expression *H. melpomene* heads, lilac expression in *H. melpomene* legs, green expression in *H. melpomene* antennae, and purple expression in *H. melpomene* mouth parts. (A) Phosphatidylinositol synthase gene family. (B) Protein C kinase 53E gene family. (C) Phospholipase D gene family. (D) Retinal degeneration A gene family. (E) Retinal degeneration B gene damily. (F) Retinal degeneration C gene family. (G) Vacuolar H+ ATPase 100kD subunit 1gene family.

**Figure S6: Phylogeny and expression of phototransduction genes 5.** Phylogenetic trees were generated using sequences from 8 insect genomes. Orange bars represent expression in *M. sexta* heads, red bars expression *H. melpomene* heads, lilac expression in *H. melpomene* legs, green expression in *H. melpomene* antennae, and purple expression in *H. melpomene* mouth parts. (A) Innexin gene family.

**Figure S7: Immunohistochemistry of an unclassified and long wavelength opsin.** Longitudinal sections of a butterfly retina. (A) Negative control for the unclassified opsin protein. For this experiment, the primary anti-UnRh was excluded only anti-LWRh and secondary antibodies were added to the sections to test for general background staining. (B) Diagonal section shows that UnRh is predominantly found in the upper 1/4^th^ of the butterfly ommatidia.

**Table S1: Accession numbers for RNA-Seq data used.**

**Table S2: Number of phototransduction genes in 8 insect genomes.**

**Table S3: *H. melpomene* BLAST results and annotations.**

**Table S4: *M. sexta* BLAST results and annotations.**

**Table S5: GenBank Accession numbers for *M. sexta* and *H. melpomene* phototransduction genes.**

**Table S6: Phylogenetic tree models.**

**Table S7: Annotated DE contigs in head vs. antennae comparison.**

**Table S8: Annotated DE contigs in head vs. legs comparison.**

**Table S9: Annotated DE contigs in head vs. mouth parts comparison.**

**Table S10: Functional enrichment of differentially expressed genes.**

**Table S11: Annotated contigs commonly upregulated in heads.**

## References

Arendt D. 2003. Evolution of eyes and photoreceptor cell types. Int. J. Dev. Biol. 47:563–571. doi: 10.1387/IJDB.14756332.

Bähner M, Sander P, Paulsen R, Huber A. 2000. The visual G protein of fly photoreceptors interacts with the PDZ domain assembled INAD signaling complex via direct binding of activated Gα(q) to phospholipase Cβ. J. Biol. Chem. 275:2901–2904. doi: 10.1074/jbc.275.4.2901.

Bao R, Friedrich M. 2009. Molecular evolution of the *Drosophila* retinome: exceptional gene gain in the higher Diptera. Mol. Biol. Evol. 26:1273–87. doi: 10.1093/molbev/msp039.

Battelle BA et al. 2016. Opsin Repertoire and Expression Patterns in Horseshoe Crabs: Evidence from the genome of *Limulus polyphemus* (Arthropoda: Chelicerata). Genome Biol. Evol. 8:1571–1589. doi: 10.1093/gbe/evw100.

Bernard GD. 1983a. Bleaching of rhabdoms in eyes of intact butterflies. Science. 219:69–71.

Bernard GD. 1983b. Dark-Processes following photoconversion of butterfly rhodopsins. Biophys Struct Mech. 9:277–286.

Bloomquist BT et al. 1988. Isolation of a putative phospholipase C gene of *Drosophila, norpA*, and its role in phototransduction. Cell. 54:723–733. doi: 10.1016/S0092-8674(88)80017-5.

Briscoe AD et al. 2010. Positive selection of a duplicated UV-sensitive visual pigment coincides with wing pigment evolution in *Heliconius* butterflies. Proc. Natl. Acad. Sci. U. S. A. 107:3628–33. doi:10.1073/pnas.0910085107.

Briscoe AD. 2008. Reconstructing the ancestral butterfly eye: focus on the opsins. J. Exp. Biol. 211:1805–13.

Brotz TM, Gundelfinger ED, Borst A. 2001. Cholinergic and GABAergic pathways in fly motion vision. BMC Neurosci. 2:1. doi: 10.1186/1471-2202-2-1.

Camacho C et al. 2009. BLAST+: architecture and applications. BMC Bioinformatics. 10:421. doi:Artn 421\nDoi 10.1186/1471-2105-10-421.

Chevesich J, Kreuz AJ, Montell C. 1997. Requirement for the PDZ domain protein, INAD, for localization of the TRP store-operated channel to a signaling complex. Neuron. 18:95–105. doi: 10.1016/S0896-6273(01)80049-0.

Chung BY, Kilman VL, Keath JR, Pitman JL, Allada R. 2009. The GABAA receptor RDL acts in peptidergic PDF neurons to promote sleep in *Drosophila*. Curr. Biol. 19:386–390. doi: 10.1016/j.cub.2009.01.040.

Chung W, Marshall NJ. 2017. Complex visual adaptations in squid for specific tasks in different environments. 8:1–16. doi:10.3389/fphys.2017.00105.

Chyb S, Raghu P, Hardie RC. 1999. Polyunsaturated fatty acids activate the *Drosophila* light-sensitive channels TRP and TRPL. Nature. 397:255–259. doi: 10.1038/16703.

Conesa A et al. 2005. Blast2GO: A universal tool for annotation, visualization and analysis in functional genomics research. Bioinformatics. 21:3674–3676. doi: 10.1093/bioinformatics/bti610.

Conesa A, Götz S. 2008. Blast2GO: A comprehensive suite for functional analysis in plant genomics. Int. J. Plant Genomics. 619832. doi:10.1155/2008/619832.

Dabney A, Storey JD. 2013. qvalue: Q-value estimation for false discovery rate control.

Davey JW et al. 2016. Major improvements to the *Heliconius melpomene* genome assembly used to confirm 10 chromosome fusion events in 6 million years of butterfly evolution. G3. 6:695– 708.

Dolph PJ et al. 1993. Arrestin function in inactivation of G protein-coupled receptor rhodopsin in vivo. Science. 260:1910–1916. doi:10.1126/science.8316831.

Duan Y et al. 2017. Transcriptome analysis of molecular mechanisms responsible for light-stress response in *Mythimna separata* (Walker). Sci. Rep. 7:45188. doi: 10.1038/srep45188.

Edgar RC. 2004. MUSCLE: Multiple sequence alignment with high accuracy and high throughput. Nucleic Acids Res. 32:1792–1797. doi:10.1093/nar/gkh340.

Faggionato D, Serb JM. 2017. Strategy to identify and test putative light-sensitive non-opsin G-protein-coupled receptors: A case study. Biol. Bull. 233:70–82. doi:10.1086/694842.

Fain GL, Hardie R, Laughlin SB. 2010. Phototransduction and the evolution of photoreceptors. Curr. Biol. 20:R114-24. doi:10.1016/j.cub.2009.12.006.

Feuda R, Marle F, Bentley MA, Holland PWH. 2016. Conservation, duplication, and divergence of five opsin genes in insect evolution. Genome Biol. Evol. 8:579–587. doi:10.5287/bod-leian.

Fichelson P, Brigui A, Pichaud F. 2012. Orthodenticle and Kruppel homolog 1 regulate *Drosophila* photoreceptor maturation. Proc. Natl. Acad. Sci. U. S. A. 109:7893–8. doi:10.1073/pnas.1120276109.

Fowler MA, Sidiropoulou K, Ozkan ED, Phillips CW, Cooper DC. 2007. Corticolimbic expression of TRPC4 and TRPC5 channels in the rodent brain. PLOS One. 2. doi:10.1371/journal.pone.0000573.

French AS, Meisner S, Liu H, Weckström M, Torkkeli PH. 2015. Transcriptome analysis and RNA interference of cockroach phototransduction indicate three opsins and suggest a major role for TRPL channels. Front. Physiol. 6:00207. doi:10.3389/fphys.2015.00207.

Frentiu FD, Bernard GD, et al. 2007. Adaptive evolution of color vision as seen through the eyes of butterflies. Proc. Natl. Acad. Sci. U. S. A. 104 Suppl:8634–8640. doi:10.1073/pnas.0701447104.

Frentiu FD, Bernard GD, Sison-Mangus MP, Van Zandt Brower A, Briscoe AD. 2007. Gene duplication is an evolutionary mechanism for expanding spectral diversity in the long-wavelength photopigments of butterflies. Mol. Biol. Evol. 24:2016–2028. doi:10.1093/molbev/msm132.

Friedrich M et al. 2011. Phototransduction and clock gene expression in the troglobiont beetle *Ptomaphagus hirtus* of Mammoth cave. J. Exp. Biol. 214:3532–41.

Futahashi R et al. 2015. Extraordinary diversity of visual opsin genes in dragonflies. Proc. Natl. Acad. Sci. 112:E1247–E1256. doi:10.1073/pnas.1424670112.

Garcia-Murillas I et al. 2006. *lazaro* encodes a lipid phosphate phosphohydrolase that regulates phosphatidylinositol turnover during *Drosophila* phototransduction. Neuron. 49:533–546. doi:10.1016/j.neuron.2006.02.001.

Gengs C et al. 2002. The target of *Drosophila* photoreceptor synaptic transmission is a histamine-gated chloride channel encoded by *ort (hclA*). J. Biol. Chem. 277:42113–42120. doi:10.1074/jbc.M207133200.

Giraldo-Calderón GI, Zanis MJ, Hill CA. 2017. Retention of duplicated long-wavelength opsins in mosquito lineages by positive selection and differential expression. BMC Evol. Biol. 17:84. doi:10.1186/s12862-017-0910-6.

Girardot F, Lasbleiz C, Monnier V, Tricoire H. 2006. Specific age related signatures in *Drosophila* body parts transcriptome. BMC Genomics. 7:69. doi:10.1186/1471-2164-7-69.

Götz S et al. 2011. B2G-FAR, a species-centered GO annotation repository. Bioinformatics. 27:919–924. doi:10.1093/bioinformatics/btr059.

Götz S et al. 2008. High-throughput functional annotation and data mining with the Blast2GO suite. Nucleic Acids Res. 36:3420–3435. doi:10.1093/nar/gkn176.

Hardie RC. 2001. Phototransduction in *Drosophila melanogaster*. J. Exp. Biol. 204:3403–3409.

Hardie RC, Juusola M. 2015. Phototransduction in *Drosophila*. Curr. Opin. Neurobiol. 34:37–45. doi:10.1016/j.conb.2015.01.008.

Hardie RC, Minke B. 1992. The *trp* gene is essential for a light-activated Ca2+ channel in *Drosophila* photoreceptors. Neuron. 8:643–651. doi:10.1016/0896-6273(92)90086-S.

Hardie RC, Raghu P. 2001. Visual transduction in *Drosophila*. Nature. 413:186–193. doi:10.1038/35093002.

Haug-Collet K et al. 1999. Cloning and characterization of a Potassium-dependent Sodium/Calcium exchanger in *Drosophila*. J. Cell Biol. 147:659–669.

Henze MJ, Dannenhauer K, Kohler M, Labhart T, Gesemann M. 2012. Opsin evolution and expression in arthropod compound eyes and ocelli: Insights from the cricket *Gryllus bimaculatus*. BMC Evol. Biol. 12:163.

Horridge GA, Giddings C, Stange G. 1972. The superposition eye of skipper butterflies. Proc. R. Soc. London. Ser. B, Biol. Sci. 182:457–495. doi:10.1098/rspb.1972.0088.

Huang DW, Sherman BT, Lempicki RA. 2009. Systematic and integrative analysis of large gene lists using DAVID bioinformatics resources. Nat. Protoc. 4:44–57. doi:10.1038/nprot.2008.211.

Ile KE, Tripathy R, Goldfinger V, Renault AD. 2012. Wunen, a *Drosophila* lipid phosphate phosphatase, is required for septate junction-mediated barrier function. Development. 139:2535–2546. doi:10.1242/dev.077289.

Kanost MR et al. 2016. Multifaceted biological insights from a draft genome sequence of the tobacco hornworm moth, *Manduca sexta*. Insect Biochem. Mol. Biol. 76:118–147. doi:10.1016/j.ibmb.2016.07.005.

Katz B, Minke B. 2009. Drosophila photoreceptors and signaling mechanisms. Front. Cell. Neurosci. 3:2. doi:10.3389/neuro.03.002.2009.

Kingston ACN, Wardill TJ, Hanlon RT, Cronin TW. 2015. An unexpected diversity of photoreceptor classes in the longfin squid, *Doryteuthis pealeii*. PLOS One. 10:1–14. doi:10.1371/journal.pone.0135381.

Kumar S, Stecher G, Tamura K. 2016. MEGA7: Molecular Evolutionary Genetics Analysis Version 7.0 for bigger datasets. Mol. Biol. Evol. 33:1870–1874. doi:10.1093/molbev/msw054.

Langmead B, Trapnell C, Pop M, Salzberg SL. 2009. Ultrafast and memory-efficient alignment of short DNA sequences to the human genome. Genome Biol. 10:R25. doi:gb-2009-10-3-r25 [pii]\n10.1186/gb-2009-10-3-r25.

Lee YJ et al. 1994. The *Drosophila dgq* gene encodes a G alpha protein that mediates phototransduction. Neuron. 13:1143–57. doi:10.1016/0896-6273(94)90052-3.

Leung H et al. 2008. DAG lipase activity is necessary for TRP channel regulation in *Drosophila* photoreceptors. Neuron. 58:884–896.

Li B, Dewey CN. 2011. RSEM: accurate transcript quantification from RNA-Seq data with or without a reference genome. BMC Bioinformatics. 12:323. doi:10.1186/1471-2105-12-323.

Li H, Durbin R. 2009. Fast and accurate short read alignment with Burrows-Wheeler transform. Bioinformatics. 25:1754–1760. doi:10.1093/bioinformatics/btp324.

von Lintig J, Kiser PD, Golczak M, Palczewski K. 2010. The biochemical and structural basis for *trans*-to-*cis* isomerization of retinoids in the chemistry of vision. Trends Biochem. Sci. 35:400–410. doi:10.1016/j.tibs.2010.01.005.

Liu CH et al. 2008. Ca2+-Dependent metarhodopsin inactivation mediated by calmodulin and NINAC myosin III. Neuron. 59:778–789. doi:10.1016/j.neuron.2008.07.007.

Liu X, Krause WC, Davis RL. 2007. GABAA receptor RDL inhibits *Drosophila* olfactory associative learning. Neuron. 56:1090–1102. doi:10.1016/j.neuron.2007.10.036.

Macias-Muñoz A, McCulloch KJ, Briscoe AD. 2017. Copy number variation and expression analysis reveals a non-orthologous *pinta* gene family member involved in butterfly vision. Genome Biol. Evol. 9:3398–3412. doi:10.1093/gbe/evx230.

Macias-Muñoz A, Smith G, Monteiro A, Briscoe AD. 2016. Transcriptome-wide differential gene expression in *Bicyclus anynana* butterflies: Female vision-related genes are more plastic. Mol. Biol. Evol. 33:79–92. doi:10.1093/molbev/msv197.

Mahato S et al. 2014. Common transcriptional mechanisms for visual photoreceptor cell differentiation among Pancrustaceans. PLOS Genet. 10:e1004484. doi:10.1371/journal.pgen.1004484.

Marygold SJ et al. 2013. FlyBase: Improvements to the bibliography. Nucleic Acids Res. 41:D751–D757. doi:10.1093/nar/gks1024.

McCulloch KJ et al. 2017. Sexual dimorphism and retinal mosaic diversification following the evolution of a violet receptor in butterflies. Mol. Biol. Evol. 34:2271–2284. doi:10.1093/molbev/msx163.

McCulloch KJ, Osorio D, Briscoe AD. 2016. Sexual dimorphism in the compound eye of *Heliconius erato*: a nymphalid butterfly with at least five spectral classes of photoreceptor. J. Exp. Biol. 219:2377–2387. doi:10.1242/jeb.136523.

Min XJ, Butler G, Storms R, Tsang A. 2005. OrfPredictor: Predicting protein-coding regions in EST-derived sequences. Nucleic Acids Res. 33:677–680. doi:10.1093/nar/gki394.

Montell C. 2012. *Drosophila* visual transduction. Trends Neurosci. 35:356–363. doi:10.1016/j.tins.2012.03.004.

Montell C. 2005. TRP channels in *Drosophila* photoreceptor cells. J. Physiol. 567:45–51. doi:10.1113/jphysiol.2005.092551.

Montell C, Birnbaumer L, Flockerzi V. 2002. The TRP channels, a remarkably functional family. Cell. 108:595–598. doi:10.1016/S0092-8674(02)00670-0.

Montell C, Rubin GM. 1989. Molecular characterization of the *Drosophila trp* locus: a putative integral membrane protein required for phototransduction. Neuron. 2:1313–1323. doi:10.1016/0896-6273(89)90069-X.

Montgomery SH, Merrill RM, Ott SR. 2016. Brain composition in *Heliconius* butterflies, posteclosion growth and experience-dependent neuropil plasticity. J. Comp. Neurol. 524:1747–1769. doi:10.1002/cne.23993.

Niemeyer BA, Suzuki E, Scott K, Jalink K, Zuker CS. 1996. The *Drosophila* light-activated conductance is composed of the two channels TRP and TRPL. Cell. 85:651–659. doi:10.1016/S0092-8674(00)81232-5.

Nilsson DE, Land MF, Howard J. 1984. Afocal apposition optics in butterfly eyes. Nature. 312:561–563. doi:10.1038/312561a0.

Okada T et al. 1998. Molecular cloning and functional characterization of a novel receptor-activated TRP Ca2+ channel from mouse brain. J. Biol. Chem. 273:10279–10287. doi:10.1074/jbc.273.17.10279.

Peng G et al. 2016. TRPA1 channels in *Drosophila* and honey bee ectoparasitic mites share heat sensitivity and temperature-related physiological functions. Front. Physiol. 7:1–10. doi:10.3389/fphys.2016.00447.

Perry M et al. 2016. Molecular logic behind the three-way stochastic choices that expand butterfly colour vision. Nature. 535:280–4. doi:10.1038/nature18616.

Pirih P, Arikawa K, Stavenga DG. 2010. An expanded set of photoreceptors in the Eastern Pale Clouded Yellow butterfly, *Colias erate*. J. Comp. Physiol. A Neuroethol. Sensory, Neural, Behav. Physiol. 196:501–517. doi:10.1007/s00359-010-0538-0.

Plachetzki DC, Degnan BM, Oakley TH. 2007. The origins of novel protein interactions during animal opsin evolution. PLOS One. 2. doi:10.1371/journal.pone.0001054.

Plachetzki DC, Fong CR, Oakley TH. 2010. The evolution of phototransduction from an ancestral cyclic nucleotide gated pathway. Proc. R. Soc. B Biol. Sci. 277:1963–1969. doi:10.1098/rspb.2009.1797.

Ploner A. 2012. Heatplus: Heatmaps with row and/or column covariates and colored clusters.

Porter ML et al. 2012. Shedding new light on opsin evolution. Proc. R. Soc. B Biol. Sci. 279:3–14. doi:10.1098/rspb.2011.1819.

Raible F et al. 2006. Opsins and clusters of sensory G-protein-coupled receptors in the sea urchin genome. Dev. Biol. 300:461–475. doi:10.1016/j.ydbio.2006.08.070.

Ramirez MD et al. 2016. The last common ancestor of most bilaterian animals possessed at least nine opsins. Genome Biol. Evol. 8:3640–3652. doi:10.1093/gbe/evw248.

Rivera AS et al. 2010. Gene duplication and the origins of morphological complexity in pancrustacean eyes, a genomic approach. BMC Evol. Biol. 10:123. doi:10.1186/1471-2148-10-123.

Robinson M, Oshlack A. 2010. A scaling normalization method for differential expression analysis of RNA-seq data. Genome Biol. 11:R25. doi:10.1186/gb-2010-11-3-r25.

Robinson MD, Mccarthy DJ, Smyth GK. 2010. edgeR?: a Bioconductor package for differential expression analysis of digital gene expression data. 26:139–140. doi:10.1093/bioinformatics/btp616.

Sakai T, Kasuya J, Kitamoto T, Aigaki T. 2009. The Drosophila TRPA channel, painless, regulates sexual receptivity in vigin females. Genes. Brain. Behav. 8:546–557. doi:10.1088/1367-2630/15/1/015008.Fluid.

Schwarz G. 1978. Estimating the dimension of a model. Ann. Stat. 6:461–464.

Seymoure BM, Mcmillan WO, Rutowski R. 2015. Peripheral eye dimensions in Longwing (*Heliconius*) butterflies vary with body size and sex but not light environment nor mimicry ring. J. Res. Lepid. 48:83–92.

Shichida Y, Matsuyama T. 2009. Evolution of opsins and phototransduction. Philos. Trans. R. Soc. B Biol. Sci. 364:2881–2895. doi:10.1098/rstb.2009.0051.

Shieh BH, Stamnes MA, Seavello S, Harris GL, Zuker CS. 1989. The ninaA gene required for visual transduction in *Drosophila* encodes a homologue of cyclosporin A-binding protein. Nature. 338:67–70. doi:10.1038/338067a0.

Shieh BH, Zhu MY. 1996. Regulation of the TRP Ca2+ channel by INAD in *Drosophila* photoreceptors. Neuron. 16:991–998. doi:10.1016/S0896-6273(00)80122-1.

Sison-Mangus MP, Briscoe AD, Zaccardi G, Knuttel H, Kelber A. 2008. The lycaenid butterfly *Polyommatus icarus* uses a duplicated blue opsin to see green. J. Exp. Biol. 211:361–369. doi:10.1242/jeb.012617.

Smith G, Chen YR, Blissard GW, Briscoe AD. 2014. Complete dosage compensation and sex-biased gene expression in the moth *Manduca sexta*. Genome Biol. Evol. 6:526–537. doi:10.1093/gbe/evu035.

Spaethe J, Briscoe AD. 2004. Early duplication and functional diversification of the opsin gene family in insects. Mol. Biol. Evol. 21:1583–1594. doi:10.1093/molbev/msh162.

Spaethe J, Briscoe AD. 2005. Molecular characterization and expression of the UV opsin in bumblebees: Three ommatidial subtypes in the retina and a new photoreceptor organ in the lamina. 2347–2361. doi:10.1242/jeb.01634.

Sperling L, Hubbard R. 1975. Squid retinochrome. J. Gen. Physiol. 65:235–251. doi:10.1085/jgp.65.2.235.

Stavenga DG, Hardie RC. 2011. Metarhodopsin control by arrestin, light-filtering screening pigments, and visual pigment turnover in invertebrate microvillar photoreceptors. J. Comp. Physiol. A Neuroethol. Sensory, Neural, Behav. Physiol. 197:227–241. doi:10.1007/s00359-010-0604-7.

Suga H, Schmid V, Gehring WJ. 2008. Evolution and functional diversity of jellyfish opsins. Curr. Biol. 18:51–55. doi:10.1016/j.cub.2007.11.059.

Terakita A, Hara R, Hara T. 1989. Retinal-binding protein as a shuttle for retinal in the rhodopsin-retinochrome system of the squid visual cells. Vision Res. 29:639–652. doi:10.1016/0042-6989(89)90026-6.

Terakita A, Yamashita T, Shichida Y. 2000. Highly conserved glutamic acid in the extracellular IV-V loop in rhodopsins acts as the counterion in retinochrome, a member of the rhodopsin family. Proc. Natl. Acad. Sci. U. S. A. 97:14263–14267. doi:10.1073/pnas.260349597.

Tsunoda S et al. 1997. A multivalent PDZ-domain protein assembles signalling complexes in a G-protein-coupled cascade. Nature. 388:243–249. doi:10.1038/40805.

Venkatachalam K et al. 2010. Dependence on a retinophilin/myosin complex for stability of PKC and INAD and termination of phototransduction. J. Neurosci. 30:11337–11345. doi:10.1523/JNEUROSCI.2709-10.2010.Dependence.

Vöcking O, Kourtesis I, Tumu S, Hausen H. 2017. Co-expression of xenopsin and rhabdomeric opsin in photoreceptors bearing microvilli and cilia. Elife. 6:e23435. doi:10.7554/eLife.23435.

Wang T et al. 2005. Light activation, adaptation, and cell survival functions of the Na +/Ca2+ exchanger CalX. Neuron. 45:367–378. doi:10.1016/j.neuron.2004.12.046.

Wang T, Wang X, Xie Q, Montell C. 2008. The SOCS box protein STOPS is required for phototransduction through its effects on phospholipase C. Neuron. 57:56–68. doi:10.1016/j.neuron.2007.11.020.

Warrant E, Dacke M. 2016. Visual navigation in nocturnal insects. Physiology. 31:182–192. doi:10.1152/physiol.00046.2015.

Wernet MF, Perry MW, Desplan C. 2015. The evolutionary diversity of insect retinal mosaics: common design principles and emerging molecular logic. Trends Genet. 31:316–328. doi:10.1016/j.tig.2015.04.006.

Wickham H. 2009. ggplot2: Elegant Graphics for Data Analysis. Springer-Verlag New York.

Xu J, Sornborger AT, Lee JK, Shen P. 2008. Drosophila TRPA channel modulates sugar-stimulated neural excitation, avoidance and social response. Nat. Neurosci. 11:676–682. doi:10.1038/nn.2119.

Xu P et al. 2013. The evolution and expression of the moth visual opsin family. PLOS One. 8:e78140. doi:10.1371/journal.pone.0078140.

Yack JE, Johnson SE, Brown SG, Warrant EJ. 2007. The eyes of *Macrosoma* sp. (Lepidoptera: Hedyloidea): A nocturnal butterfly with superposition optics.Arthropod Struct. Dev. 36:11–22. doi:10.1016/j.asd.2006.07.001.

Yagi N, Koyama N. 1963. The compound eye of Lepidoptera: Approach from organic evolution. In: Shinkyo-Press.

Yuan F, Bernard GD, Le J, Briscoe AD. 2010. Contrasting modes of evolution of the visual pigments in *Heliconius* butterflies. Mol. Biol. Evol. 27:2392–2405. doi:10.1093/molbev/msq124.

Yuan Q, Song Y, Yang C-H, Jan LY, Jan YN. 2013. Female contact modulates male aggression via a sexually dimorphic GABAergic circuit in *Drosophila*. Nat. Neurosci. 17:81–88. doi:10.1038/nn.3581.

Zhan S, Merlin C, Boore JL, Reppert SM. 2011. The monarch butterfly genome yields insights into long-distance migration. Cell. 147:1171–1185. doi:10.1016/j.cell.2011.09.052.

Zhang N, Zhang J, Purcell KJ, Cheng Y, Howard K. 1997. The *Drosophila* protein wunen repels migrating germ cells. Nature. 385:64–67. doi:10.1038/385064a0.

