## Supplementary figures and images for "Evolution of Phototransduction Genes in Lepidoptera"

### Supplemental Figure 1

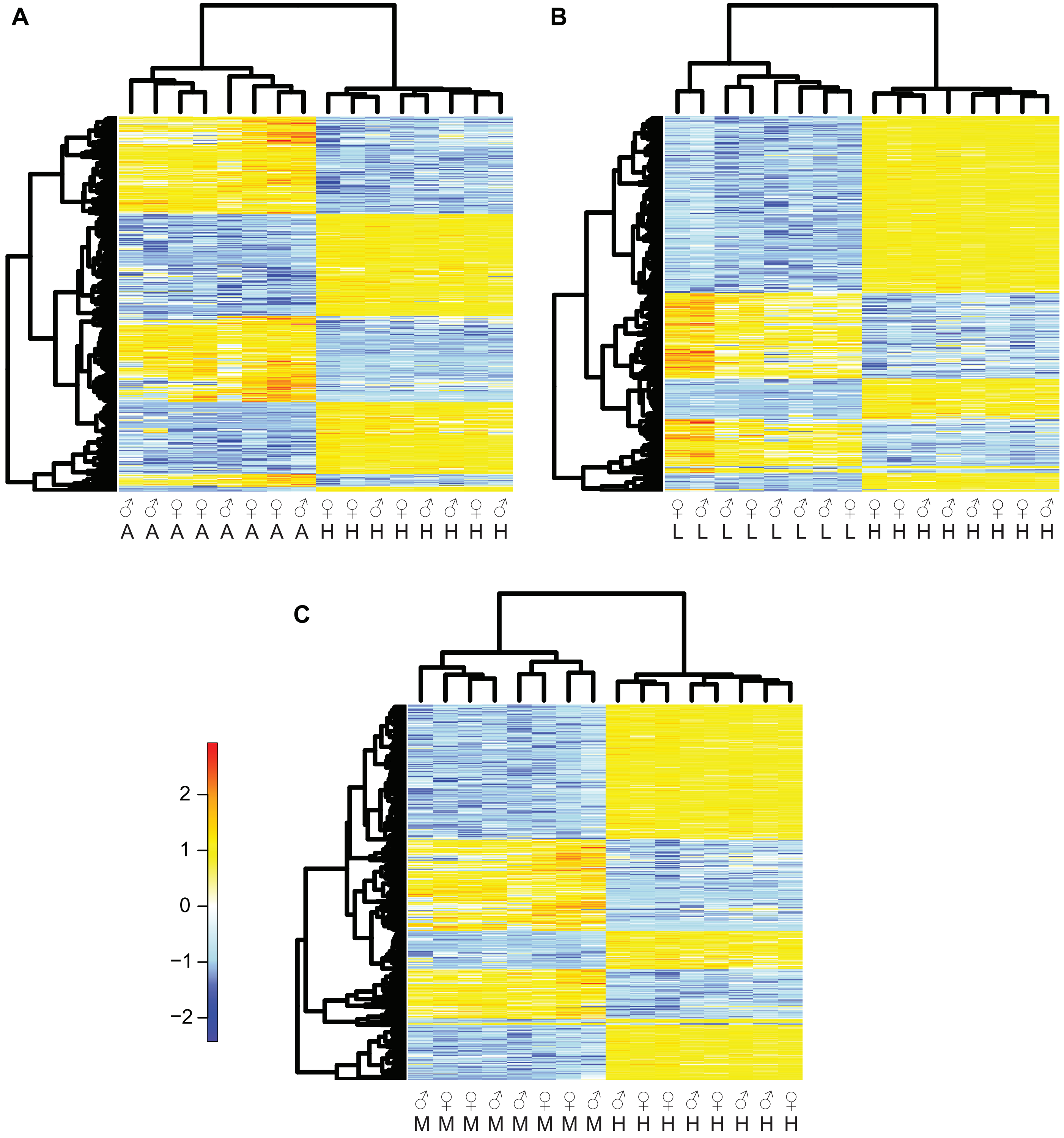

### Supplemental Figure 2

#### A. Arrestin

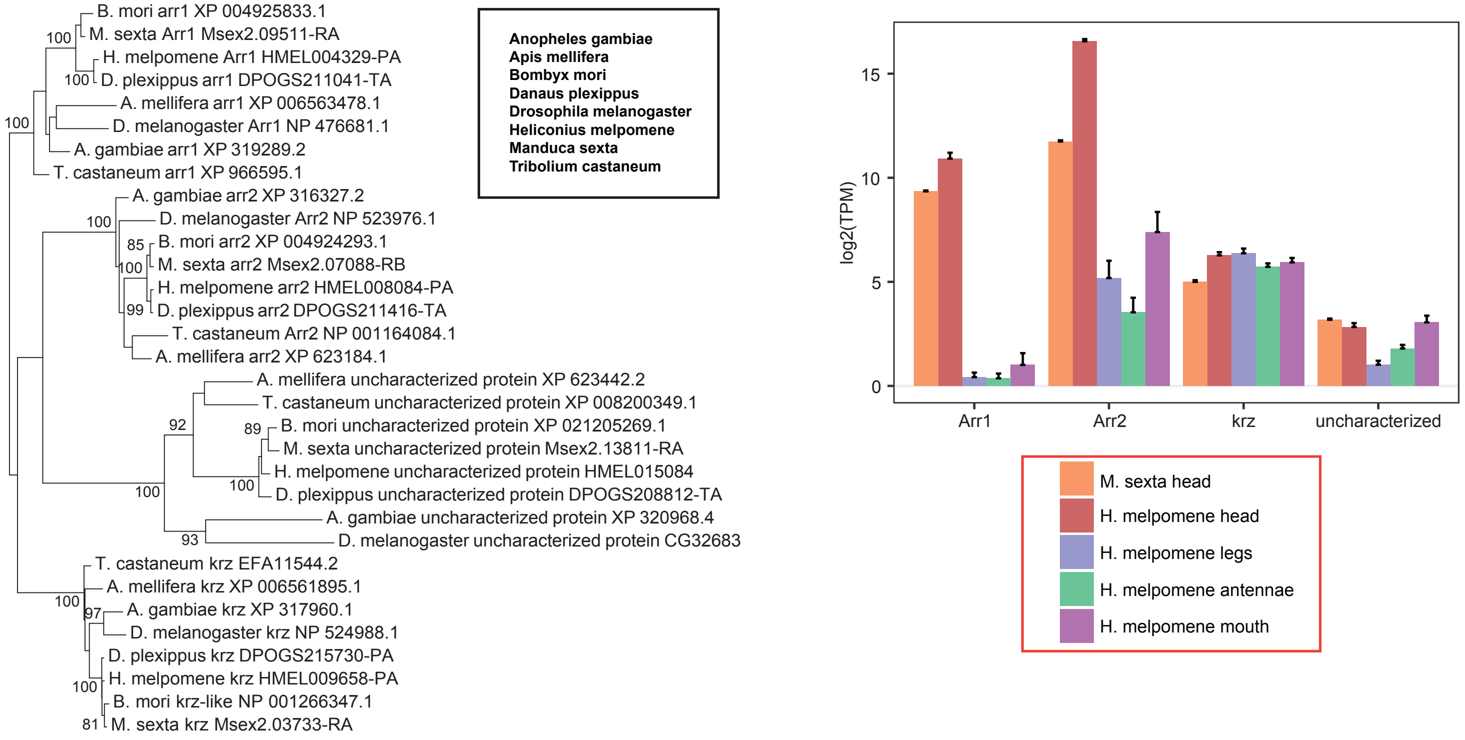


#### B. Cacophony

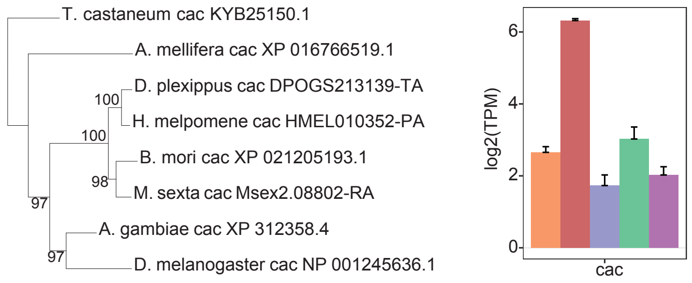


#### C. Calx Na/Ca-exchange protein

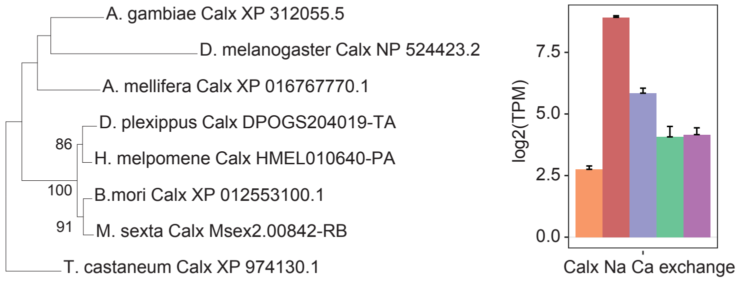


#### D. Calmodulin

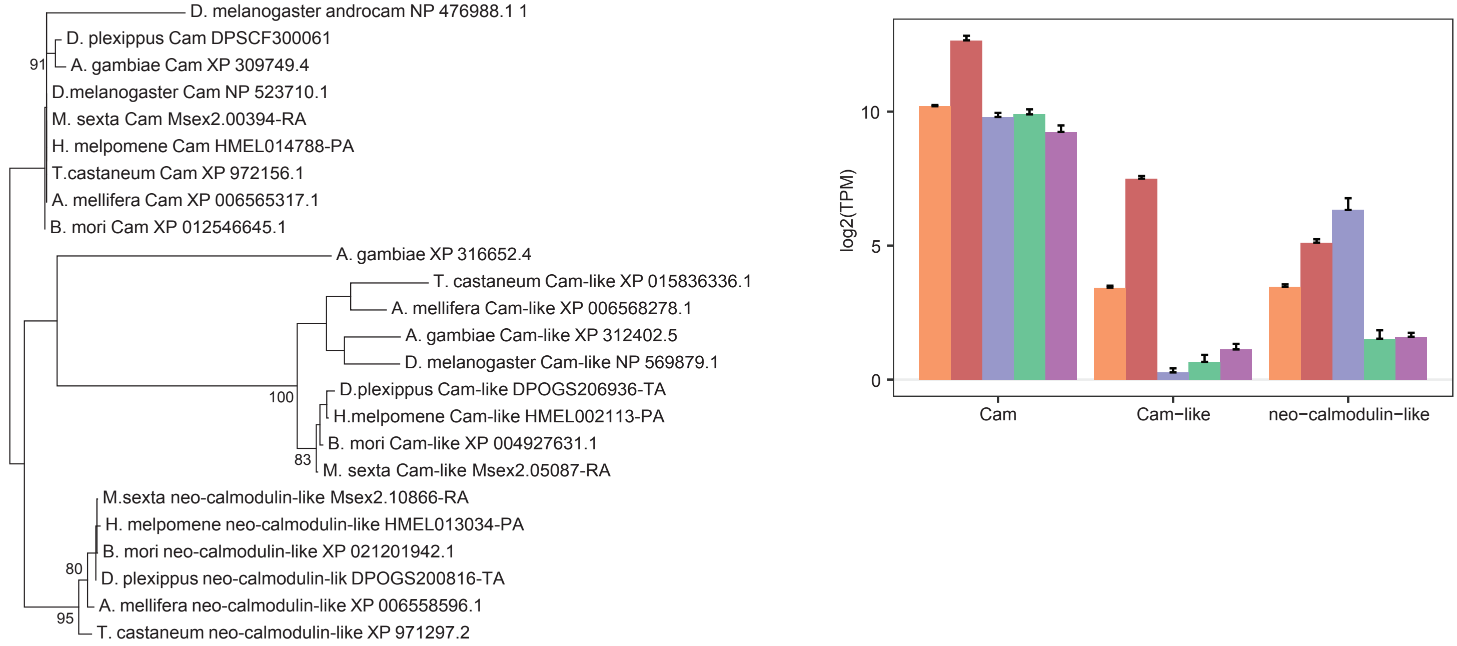

### Supplemental Figure 3

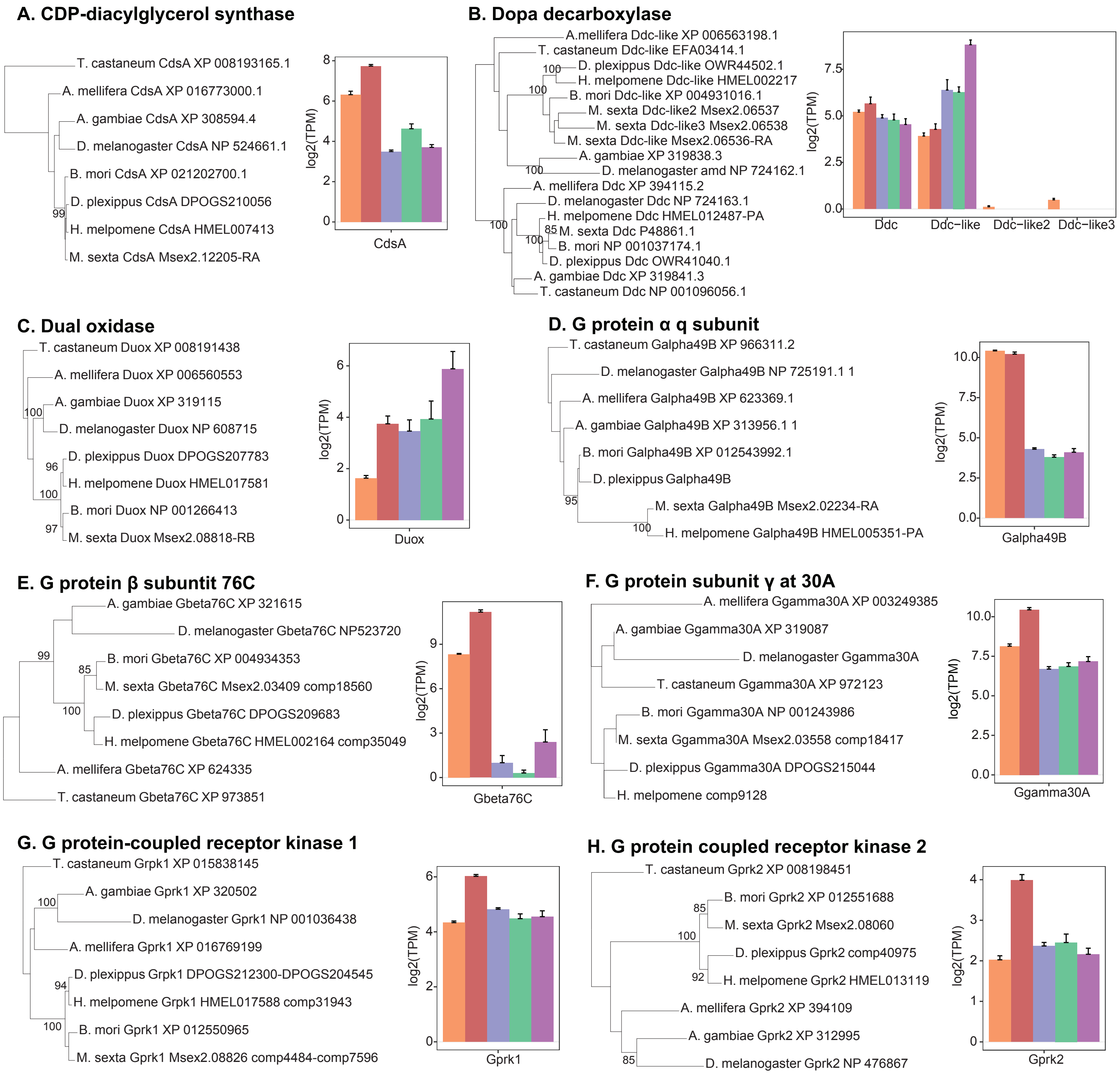

### Supplemental Figure 4

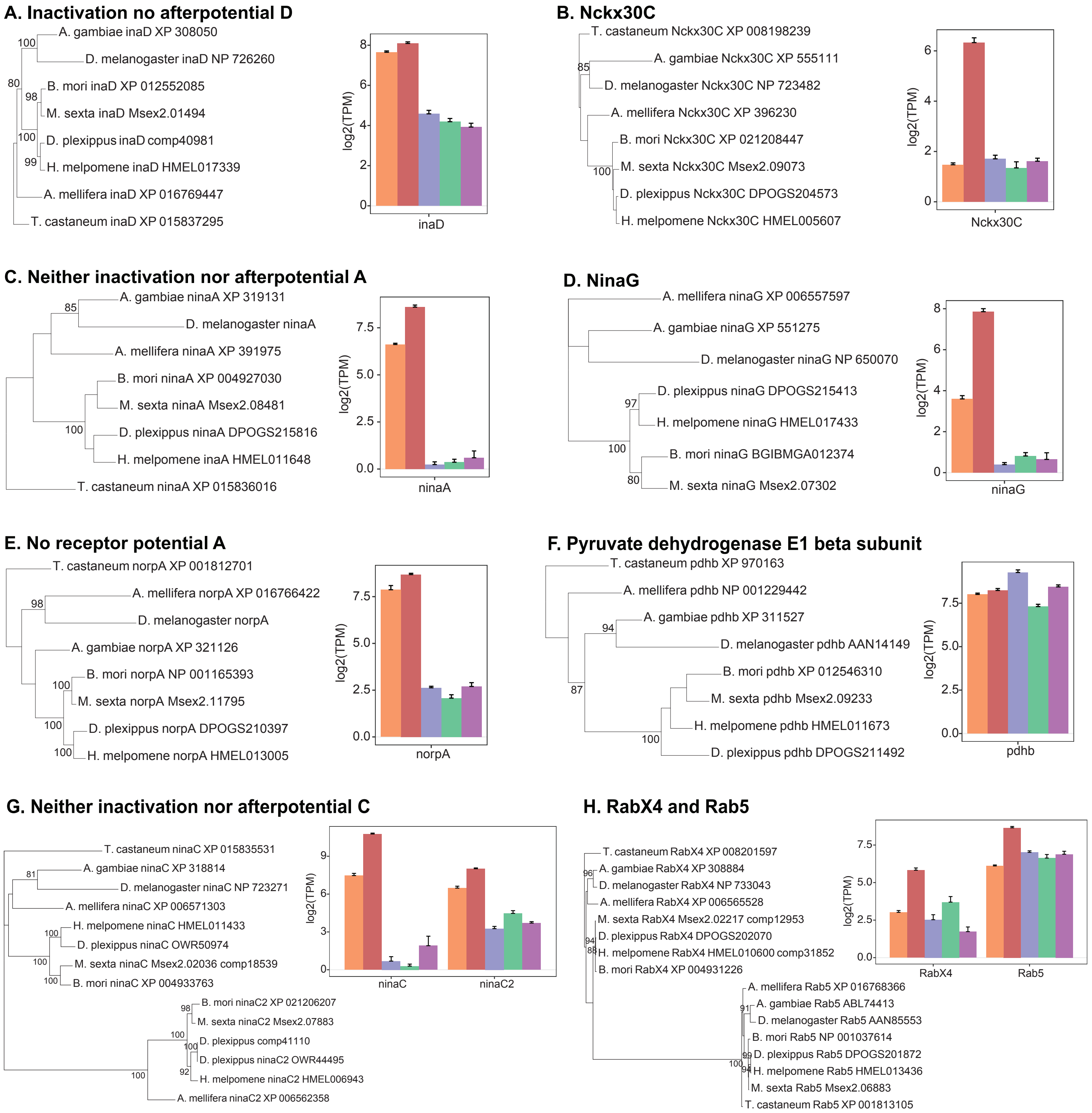

### Supplemental Figure 5

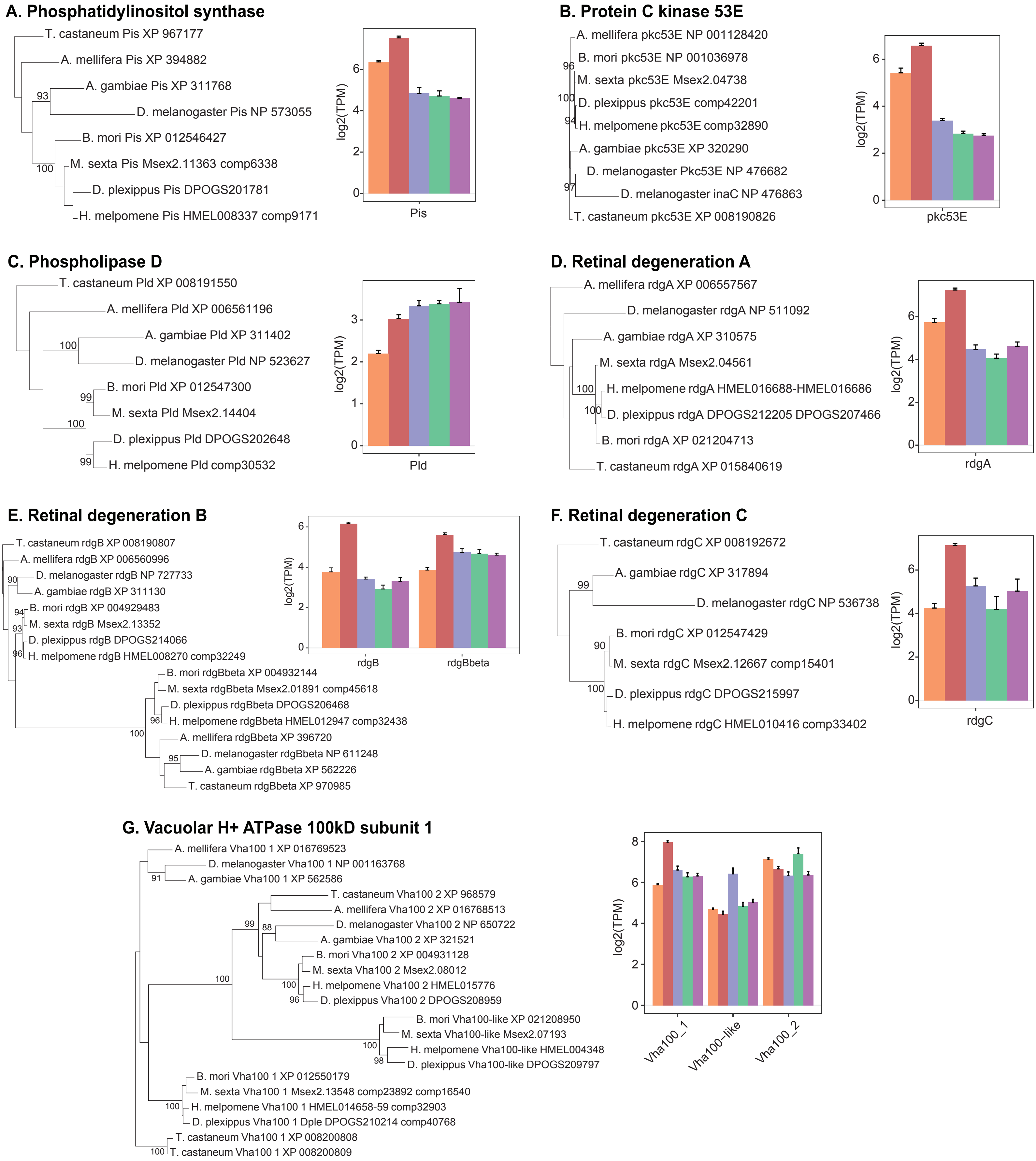

### Supplemental Figure 6

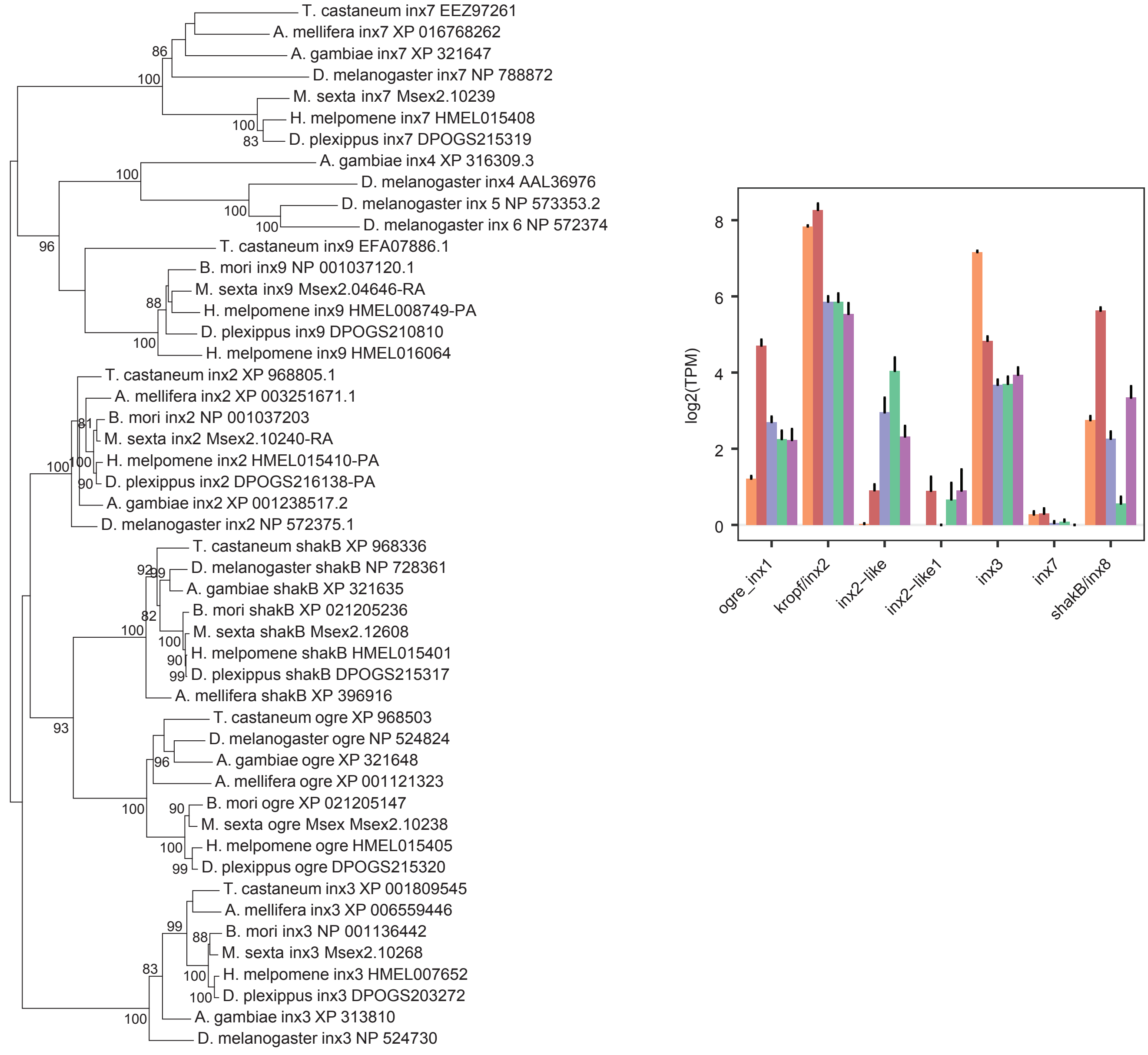

### Supplemental Figure 7

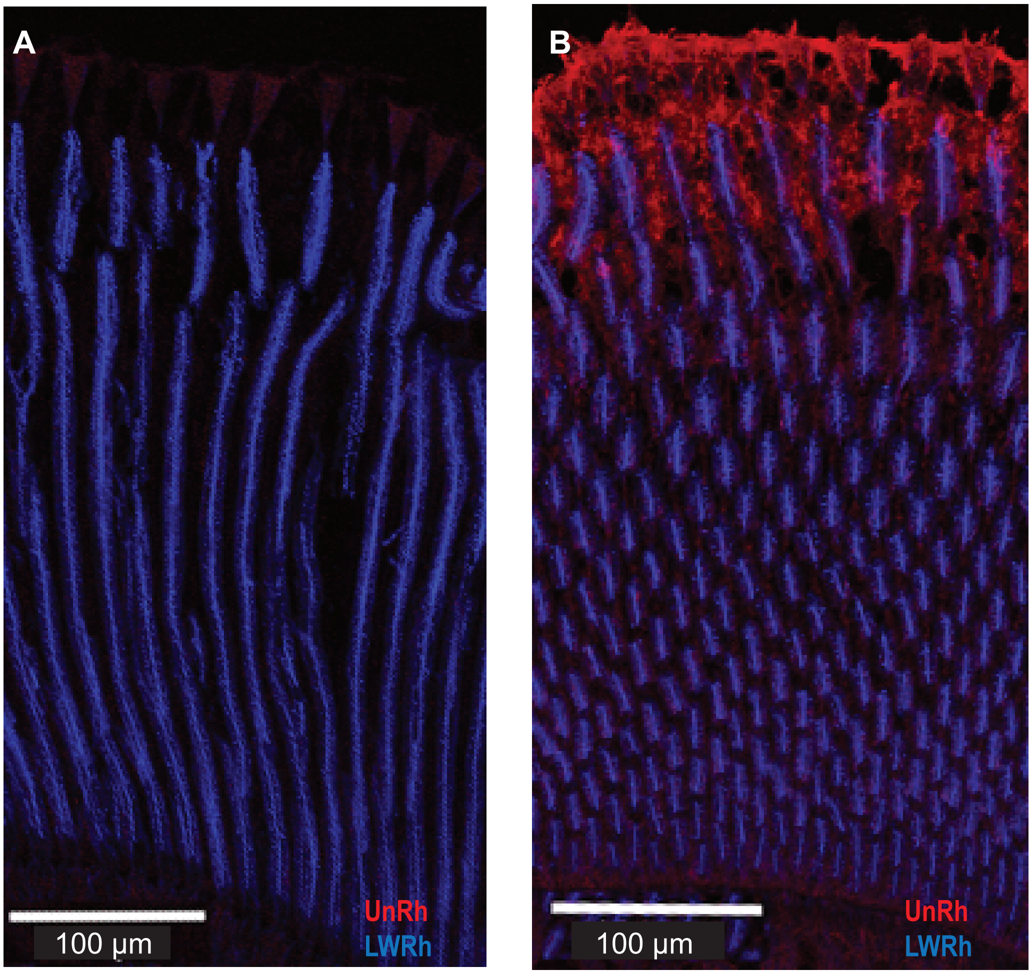
